# Modelling spinal locomotor circuits for movements in developing zebrafish

**DOI:** 10.1101/2021.02.10.430671

**Authors:** Yann Roussel, Stephanie F. Gaudreau, Emily R. Kacer, Mohini Sengupta, Tuan V. Bui

## Abstract

Many spinal circuits dedicated to locomotor control have been identified in the developing zebrafish. How these circuits operate together to generate the various swimming movements during development remains to be clarified. In this study, we iteratively built models of developing zebrafish spinal circuits coupled to simplified musculoskeletal models that reproduce coiling and swimming movements. The neurons of the models were based upon morphologically or genetically identified populations in the developing zebrafish spinal cord. We simulated intact spinal circuits as well as circuits with silenced neurons or altered synaptic transmission to better understand the role of specific spinal neurons. Analysis of firing patterns and phase relationships helped identify possible mechanisms underlying the locomotor movements of developing zebrafish. Notably, our simulations demonstrated how the site and the operation of rhythm generation could transition between coiling and swimming. The simulations also underlined the importance of contralateral excitation to multiple tail beats. They allowed us to estimate the sensitivity of spinal locomotor networks to motor command amplitude, synaptic weights, length of ascending and descending axons, and firing behaviour. These models will serve as valuable tools to test and further understand the operation of spinal circuits for locomotion.

## INTRODUCTION

Movements made in the early stages of development can be critical for the survival of many species. The escape response seen in various fish and amphibians is one such example of a vital movement present at early developmental stages (Domenici & Hale, 2019). However, the nervous system’s control of movement does not come fully formed but matures as the nervous system develops (Favero et al., 2014). This maturation enables a broader repertoire of movements to arise. During this process, new neurons are born and subsequently integrated into neural circuits that are newly formed or refined, presumably leading to the emergence of progressively more coordinated and skillful maneuvers. Determining how the assembly of new circuits leads to the emergence of new movements can provide valuable insights into the role of distinct neurons or circuits in motor control.

The maturation of swimming in developing zebrafish has been well described at both the ethological and the cellular levels (Drapeau et al., 2002; McLean & Fetcho, 2009). Single strong body bends on one side of the body, also known as coils, emerge during the first day of development at around 17 hours post-fertilization (hpf) as the earliest locomotor behaviour (Saint-Amant & Drapeau, 1998). Single coils are quickly followed by double coils (i.e. two successive coils, one for each side of the body) at around 24 hpf (Knogler et al., 2014). Touch-evoked swimming appears around 27 hpf as coiling begins to subside. Spontaneous swimming movements emerge around 2-3 days post fertilization (Saint-Amant, 2010). The first swimming movement zebrafish exhibit is burst swimming characterized by long (1 s long) but infrequent episodes of tail beats. Burst swimming is then replaced by beat-and-glide swimming characterized by shorter (several hundreds of ms long) but more frequent episodes. In both cases, swim episodes consist of repetitive left-right alternating, low-amplitude tail beats that propagate from the rostral toward the caudal end of the fish body and are generated at 20 to 80 Hz (Budick & O’Malley, 2000; Buss & Drapeau, 2001).

During this rapid series of transitions between locomotor maneuvers, populations of spinal neurons are progressively generated, starting with primary motoneurons at about 9 hpf. Subsequently, spinal motoneurons and interneurons are generated in stereotyped spatiotemporal birth orders (Kimmel et al., 1994; Myers et al., 1986; Satou et al., 2012). Two successive waves of axogenesis occur in the embryonic spinal cord (Bernhardt et al., 1990). The first wave occurs around 16-17 hpf. It includes axon growth in primary motoneurons (MNs) that innervate red and white muscle fibres at early developmental stages (Buss & Drapeau, 2000). Primary MNs enable coiling and escape movements (Kimmel et al., 1995; Saint-Amant & Drapeau, 2000). Several spinal interneurons that are also important for early movements extend their axons along with primary MNs. These include Ipsilateral Caudal (IC) interneurons that are thought to play an essential role in driving the rhythm of early locomotor behaviour due to their endogenous bursting activity (Tong & McDearmid, 2012). The second wave of axon growth occurs at around 23-25 hpf. It involves axon growth in secondary motoneurons involved with slower movements (D. W. Liu & Westerfield, 1988) and spinal interneuron populations that include excitatory and inhibitory, ipsilaterally and contralaterally, and ascending and descending projecting subtypes (Bernhardt et al., 1990; Higashijima et al., 2004). The progressive generation of new populations of spinal neurons and continued axonal growth coincides with the expansion of the zebrafish locomotor repertoire. This timing suggests that incorporating spinal circuits into existing locomotor circuits underlies the acquisition of novel locomotor maneuvers.

We have recently provided evidence that the maturation from coiling to later stages of swimming is accompanied by an operational switch in how spinal locomotor circuits generate the rhythm underlying tail beats. Specifically, we demonstrated that spinal circuits transitioned from relying upon pacemakers with endogenous bursting properties during coiling towards depending upon network oscillators whose rhythm is driven by excitatory and inhibitory synapses (Roussel et al., 2020). In light of these and earlier findings describing the composition and maturation of spinal locomotor circuits, we sought to generate computational models that replicate developmental locomotor movements of the zebrafish. We iteratively constructed models for several locomotor movements by incorporating specific spinal populations, shifts in relative connection strength, and changes in the firing behaviour of neurons. While computational modelling has generated invaluable insights into the function and mechanisms of spinal locomotor circuits of several species (Ausborn et al., 2019; Bicanski et al., 2013; Danner et al., 2019; Ferrario et al., 2018; Hull et al., 2016; A. K. Kozlov et al., 2014; Sautois et al., 2007), there is to our knowledge no such model for the developing zebrafish spinal cord. Here, we build some of the first computational models of the zebrafish spinal locomotor circuit that can accurately reproduce predominant locomotor behaviours during early zebrafish development. In the process, we test theories about the possible contributions of specific neural circuits and spinal populations to locomotor movements in zebrafish and identify untested hypotheses on the operation of spinal locomotor networks in developing zebrafish.

## RESULTS

We aimed to model how new locomotor movements may emerge from the integration of spinal interneurons and the modification of synaptic weights and firing behaviour over the first few days of development in the zebrafish. Our approach was to build an initial model based upon previously reported experimental observations of spinal circuits when the first locomotor movements emerge in zebrafish around 1 dpf. We then successively built upon this initial model to replicate several locomotor maneuvers of the developing zebrafish.

The models were composed of single-compartment neurons whose firing dynamics were determined by a small set of differential equations (Izhikevich, 2007). The firing of motoneurons was converted to muscle output. This output was used to estimate body angle and locomotor activity during simulations (**Figure 1**). The composition of each model depended on the developmental stage and the locomotor movement to be generated.

**Figure 1.**
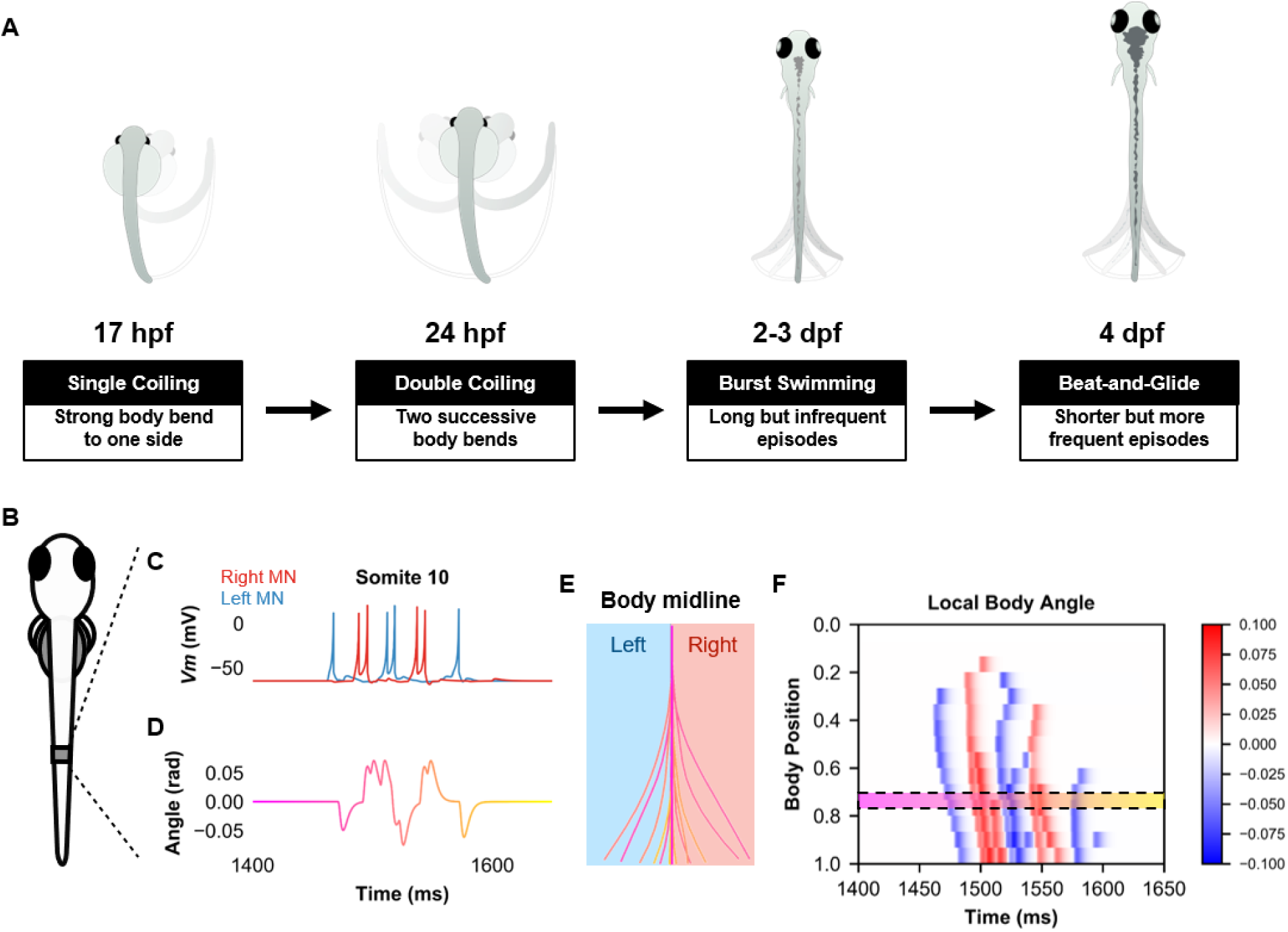
Simulation of the spinal locomotor circuit coupled to a musculoskeletal model during a beat-and-glide swimming episode. **(A)** Schematic of locomotor movements during the development of zebrafish. (**B**) Schematic of a fish body with 10^th^ somite outlined. (**C**) Motoneuron membrane potential (*Vm*) in the 10^th^ somite during a single beat-and-glide swimming episode from our model is used to calculate this body segment’s body angle variation (**D**) in a musculoskeletal model. (**E**) Several representative body midlines from this episode of beat-and-glide swimming. Body midline is computed by compiling all the calculated local body angles along the simulated fish body. (**F**) Heat-map of local body angle (in radians) across the total body length and through time during the episode. Red is for right curvatures, while blue labels left curvatures. Body position on the ordinate, 0 is the rostral extremity, while 1 is the caudal extremity. In **D**-**F**, the magenta to yellow color coding represents the progression through the swimming episode depicted.

### Single coiling results from unilateral gap junction coupling

Coiling, which is already observed at 1 dpf, is characterized by a single strong, slow (hundreds of ms in duration) tail beat on one side of the body followed by a return to resting position (Saint-Amant & Drapeau, 1998). Coiling events are relatively infrequent, reaching a maximum frequency of 1 Hz around 20 hpf (Saint-Amant & Drapeau, 1998). Previous studies have established that this behaviour is generated by a spinal circuit relying primarily on gap junctions (i.e. electrical synapses) (Saint-Amant & Drapeau, 2001). It has been proposed that rostrally-located IC pacemaker spinal neurons (Tong & McDearmid, 2012) drive periodic depolarizations of ipsilateral MNs via electrical synapses (Drapeau et al., 2002; Saint-Amant & Drapeau, 2001). Glycinergic synaptic bursts are observed in MNs during contralateral coiling events (Saint-Amant & Drapeau, 2001). These synaptic bursts have been proposed to arise from contralaterally projecting glycinergic neurons (Saint-Amant & Drapeau, 2000) but are not responsible for any action potential firings or coiling movements (Saint-Amant & Drapeau, 2001). Applying a gap junction blocker, heptanol, but not glutamatergic and glycinergic antagonists, suppressed spinal activity responsible for coiling (Saint-Amant & Drapeau, 2000).

NETWORK DESCRIPTION (**Figure 2*A***) Based on the experimental observations reported above, the model for single coiling consisted of rostrocaudal chains of electrically coupled spinal neurons driven by a kernel of five recurrently connected pacemakers (IC neurons). One chain consisted of ten MNs. The other chain consisted of ten contralaterally projecting commissural inhibitory neurons. Neurons from the V0d population are active during large amplitude movements such as escapes (Satou et al., 2020), and so we assumed V0ds were the commissural inhibitory neurons active during coiling, which is another large amplitude movement. We selected an IC kernel size of five as a trade-off between computational simplicity and robustness of the kernel to the failure of firing of a small number of cells. Similarly, the size of the coiling model was set to ten somites. Thus, each model somite represents approximately three biological somites. This choice was made as a trade-off between computational simplicity and recreating the kinematics of coiling fish.

**Figure 2.**
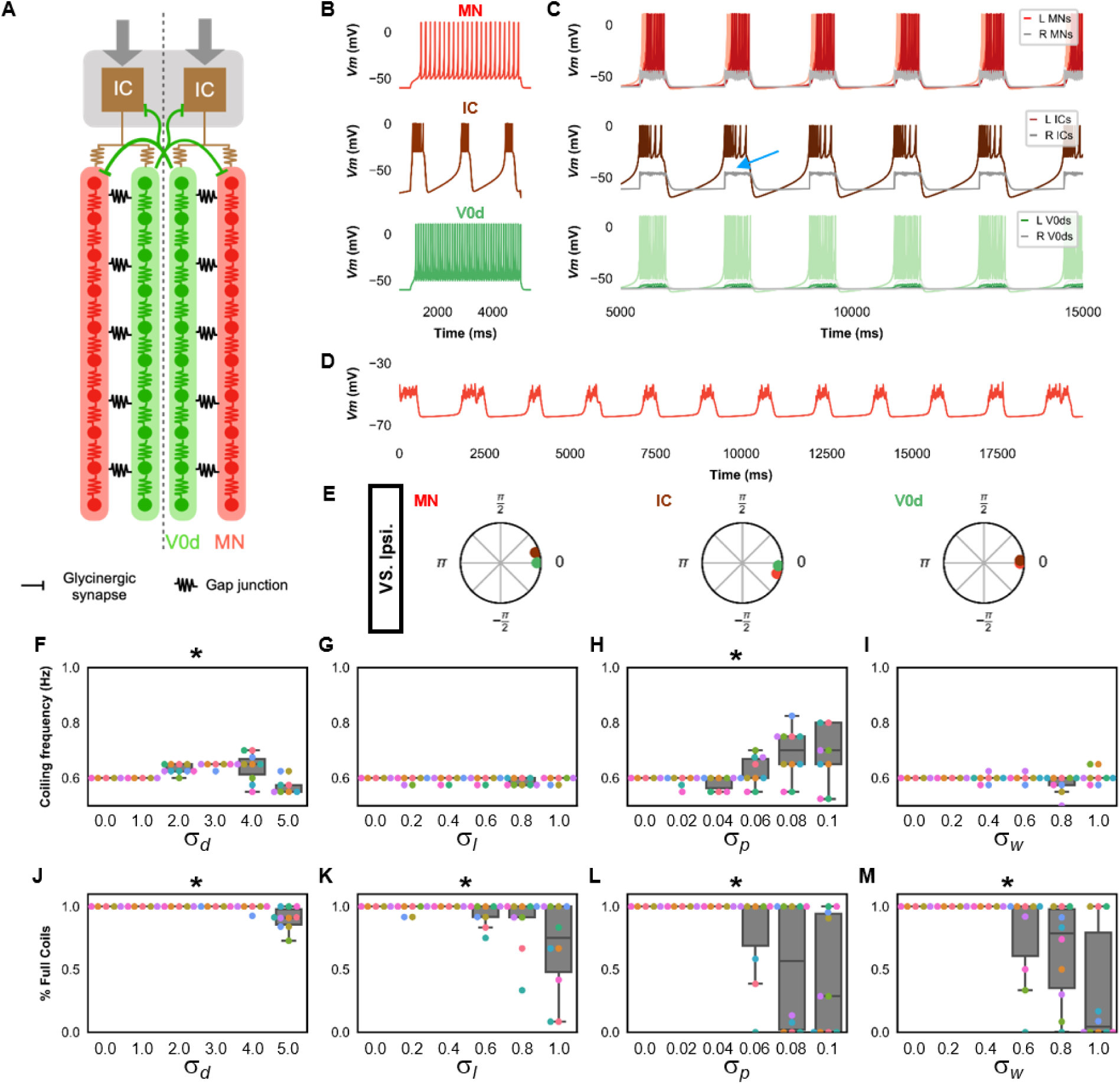
Single coiling model driven by pacemaker neurons. (**A**) Schematic of the single coiling model. The dashed line indicates the body midline. Gray arrows indicate descending motor command. (**B**) Membrane potential (*Vm*) response of isolated spinal neuron models to a depolarizing current step. (**C**) *Vm* of spinal neurons during a simulation with a tonic command to left pacemakers only. Note the synaptic bursts in grey in the right MNs and IC neurons (a blue arrow marks an example). The *Vm* of a rostral (lightest), middle, and caudal (darkest) neuron is shown, except for IC neurons that are all in a rostral kernel. (**D**) Periodic depolarizations in a hyperpolarized motoneuron on the same side where single coils are generated. (**E**) The phase delay of left neurons in relation to ipsilateral spinal neurons in the 1st somite and an IC in the rostral kernel in a 10,000 ms simulation. The reference neuron for each polar plot is labelled, and all neurons follow the same color-coding as the rest of the figure. A negative phase delay indicates that the reference neuron precedes the neuron to which it is compared. A phase of 0 indicates that a pair of neurons is in-phase; a phase of π indicates that a pair of neurons is out-of-phase. Sensitivity testing showing (**F-I**) coiling frequency and (**J-M**) proportion of full coils during ten 20,000 ms simulation runs at each value of *σ*_*d*_, *σ*_*l*_, *σ*_*p*_, and *σ*_*w*_ tested. Each run is color-coded. L: left, R: right. ***Statistics***: Asterisks denote significant differences detected using a one-factor ANOVA test. (**F**) *F*_5,59_ = 10.4, p = 5.2 x 10^-7^. (**G**) *F*_5,59_ = 2.4, p = 0.05. (**H**) *F*_5,59_ = 5.2, p = 0.0006. (**I**) *F*_5,59_ = 2.2, p = 0.07. (**J**) *F*_5,59_ = 10.9, p = 2.7 x 10^-7^. (**K**) *F*_5,59_ = 4.9, p = 0.0009. (Note that there were no pairwise differences detected). (**L**) *F*_5,59_ = 6.5, p = 8.2 x 10^-5^. (**M**) *F*_5,59_ = 8.8, p = 3.5 x 10^-6^. P-values for t-tests are found in ***Figure 2* – *source data 1***. See also ***Figure 2* -*figure supplement 1*** and ***2*** and ***Figure 2* - *video 1*** and ***2*.**

IC neurons have been reported to project caudally through multiple somites (Bernhardt et al., 1990). Therefore, in addition to their recurrent connections, each IC formed electrical synapses with several rostral MNs and V0ds (the first four of each ipsilateral chain in our model). Electrical coupling between many populations of early-born spinal neurons has been previously demonstrated, including between IC and motoneurons (Saint-Amant & Drapeau, 1998). Coupling between IC neurons and commissural inhibitory neurons has not been demonstrated yet. We based this electrical coupling between ICs and V0ds on the fact that glutamatergic blockers do not block glycinergic synaptic bursts present at this stage (Saint-Amant & Drapeau, 2001), suggesting that gap junctions mediate the activation of V0ds underlying these glycinergic bursts. Gap junction weights are found in **Table 3** in Material and Methods.

The connectivity within the chains was identical for both MN and V0d chains. Each neuron in a chain formed electrical synapses with its three nearest rostral and caudal neighbours within the same chain. There was also electrical coupling across the two ipsilateral chains as MNs formed gap junctions with the three nearest rostral and three nearest caudal V0ds and vice-versa. Paired recordings of MNs and V0ds at this stage have yet to be published. Our assumption that MNs and V0ds are electrically coupled at this stage was based upon the widespread electrical coupling between ipsilateral spinal neurons (Saint-Amant & Drapeau, 2001). To reproduce the glycinergic bursts observed in MNs at this stage (Saint-Amant & Drapeau, 2001), V0ds projected to contralateral MNs. Thus, V0ds formed glycinergic synapses with contralateral MNs and ICs. The reversal potential of glycinergic synapses is depolarized during development (Ben-Ari, 2002) and was set to -45 mV in the single coiling model (see **Table 5** in Material and Methods). All V0ds sent ascending projections to contralateral ICs. V0ds projected to contralateral MNs within five to six segments so that the *i^th^* V0d projected to all contralateral MNs between the *i-5* and *i+5* segments. Chemical synaptic weights are found in **Table 4** in Material and Methods.

Each neuron was modelled as a single compartment neuron with subthreshold and suprathreshold membrane potential dynamics described by a small set of differential equations (Izhikevich, 2007). These equations have nine parameters: *a, b, c*, *d,* and *V_max_* (which respectively represent the time scale of the recovery variable *u*, the sensitivity of *u* to the sub-threshold variation of *V*, the reset value of *V* after a spike, the reset value of *u*, and the action potential peak), and *k*, *C*, *V_r_*, and *V_t_* (coefficient for the approximation of the subthreshold part of the fast component of the current-voltage relationship of the neuron, cell capacitance, resting membrane potential, and threshold of action potential firing). Parameter values of ICs (see **Table 2** in Material and Methods for all neuron parameters) were chosen such that they exhibited a relatively depolarized threshold of action potentials and bursts of short action potentials lasting hundreds of ms as seen in experimental recordings in embryonic zebrafish (Tong & McDearmid, 2012). They were also modelled to exhibit periodic bursts lasting hundreds of ms in response to a constant tonic drive (**Figure 2*B***). This firing pattern was generated in part by having a low value of *a* and a relatively depolarized value of *c*. MNs (Drapeau et al., 1999) and V0ds were modelled to generate tonic repetitive firing in response to a step depolarization (**Figure 2*B***). Finally, to activate the circuit, a constant drive was provided to the left ICs only. Restricting the drive to left ICs prevented the appearance of near-coincident bilateral coils that could be misinterpreted as spinally mediated multiple coils.

SIMULATION RESULTS Our simulations show that this model can generate single coils characterized by large body bends to one side of the body lasting approximately one second (**Figure 2*C***, **Figure 2 - video 1**). Our base single coiling model generated six evenly interspersed single coils during a 10 s simulation. This 0.6 Hz coiling frequency is within the 0-1.0 Hz range of frequencies observed during zebrafish development (Saint-Amant & Drapeau, 1998, 2000). Silencing ICs blocked activity in all spinal neurons (**Figure 2 – figure supplement 1*A***), emphasizing the central role of the IC kernel in the generation of single coils.

Previously reported whole-cell patch-clamp recordings of MNs at this developmental stage display two types of events (Saint-Amant & Drapeau, 2000, 2001): periodic depolarizations (PDs) via electrical synapses and synaptic bursts (SBs) from contralateral spinal glycinergic neurons that are depolarizing at rest due to the depolarized chloride reversal potential observed early in development. These events last hundreds of ms. In our model, SBs were observed in the contralateral ICs and MNs (events during coilings in left neurons seen in the grey traces in **Figure 2*C***). SBs were caused by glycinergic input from V0ds activated during the ipsilateral coilings. As observed experimentally (Saint-Amant & Drapeau, 2001), preventing SBs by silencing glycinergic synapses from V0ds did not preclude the generation of single coiling, nor did it lead to the generation of multiple coilings (**Figure 2 - figure supplement 1*C***). PDs can be unmasked by hyperpolarizing motoneurons sufficiently to prevent the firing of action potentials (**Figure 2*D***). An analysis of the phase delays between ipsilateral neurons during single coils shows that IC neuron firing precedes ipsilateral MN and V0d firing (**Figure 2*E***) and reinforces that ICs drive single coiling events.

To further validate the model, we tested whether the model could still generate single coils with different parameters. First, we tested whether the model could still generate single coils when the number of model somites was increased from ten to thirty to be closer to the number of biological somites in zebrafish (Stickney et al., 2000). A thirty-somite model with IC axons extending to all somites and several modified gap junction weights (**Table 3**) generated single coils (**Figure 2 - figure supplement 2**).

Next, the base model’s sensitivity to within-model parameter variability was tested. Variability in the amplitude of the tonic motor command, the rostrocaudal extent of every axonal projection, every parameter that set the dynamics of the membrane potential of each neuron (*a*, *b*, *c*, *d*, and *V*_max_, *k*, *C*, *V_r_* and *V_t_*), and all of the weights of gap junction and chemical synapses were modelled by scaling each value by a random number picked for each simulation. The random numbers were derived from a Gaussian distribution with mean, µ = 1, and standard deviations, *σ*_*d*_ (tonic drive), *σ*_*l*_ (rostrocaudal length of axonal projections), *σ*_*p*_ (dynamics of membrane potential), and *σ*_*w*_ (synaptic weights), respectively. Ten 20-s long simulations were run at various values of *σ*_*d*_, *σ*_*l*_, *σ*_*p*_, and *σ*_*w*_. In each simulation, the variability of only one of the four sets of parameters (amplitude of motor drive, length of axonal projection, membrane potential dynamics, synaptic weights) was tested, and the standard deviations of the three other sets of parameters were set to 0.

The single coiling model’s suitability was assessed by the relative absence of truncated coils, which were movements with only partial contractions restricted to the body’s rostral segments (**Figure 2 video 2**). We sought to determine the upper limit of variability within which the single coiling model remained suitable. For this reason, the ranges of *σ*_*d*_, *σ*_*l*_, *σ*_*p*_, and *σ*_*w*_ that were tested differed amongst the four sets of parameters tested (**Figure 2*F*-*M***). A comparison of the level of variability at which the models start generating more varying frequency of coiling and more truncated coils suggests that the single coiling model is more robust to noise in the amplitude of the tonic motor command (**Figure 2*F*,*J***) and was most sensitive to variability in the parameters governing the dynamics of the membrane potential (**Figure 2*H*,*L***). The single coiling model was relatively mildly sensitive to variability in the synaptic weights and the rostrocaudal extent of the axon projections (**Figure 2*G,I,K,M***).

Overall, the model replicated this first locomotor behaviour of zebrafish in terms of the duration and frequency of coiling events as well as synaptic events of motoneurons. We then built upon this model to replicate the next step in the development of locomotion: the appearance of double coiling.

### Double coiling depends on the timing and strength of contralateral excitation and inhibition

After single coils appear, double coils emerge as a transitory locomotor behaviour at around 24 hpf, coexisting with the single coiling behaviour (Knogler et al., 2014). Double coiling is characterized by two successive coils, one on each side of the body, and lasts about one second (Knogler et al., 2014). Eventually, double coiling becomes the predominant coiling behaviour. Double coiling can represent nearly three-quarters of all coiling events at its peak frequency, with the rest mainly being single coils (Knogler et al., 2014).

At the stage when double coiling appears (24 dpf), the previous electrical scaffold for single coils seems to be supplemented with chemical glutamatergic synapses to form a hybrid electrical-chemical circuit (Knogler et al., 2014). Blocking glutamatergic transmission precludes double coils while sparing single coils (Knogler et al., 2014). In contrast, blocking glycinergic synapses led to triple or even quadruple coils (Knogler et al., 2014). These experimental observations suggest that synaptic excitation is required for successive coils after a first coil. Glycinergic transmission seems to prevent the generation of more than two successive coils. Patch-clamp recordings of MNs at this developmental stage exhibit the same isolated PDs and SBs from earlier developmental stages and show mixed events in which a PD event immediately follows an SB or vice-versa (Knogler et al., 2014). Interestingly, the application of CNQX eliminates mixed PD-SB events but not single isolated SBs, suggesting that the coupling of PD and SB in mixed events is glutamatergic (Knogler et al., 2014). Therefore, we aimed to generate a model with the following characteristics: 1) double coils lasting about one second in duration accounting for over half of the coiling events, 2) a dependence of double coiling upon excitatory synaptic transmission, 3) an increase in multiple coiling events in the absence of inhibitory synaptic transmission, and 4) the presence of mixed PD-SB events with similar sensitivity to the blockade of excitatory synaptic transmission as double coils.

NETWORK DESCRIPTION (**Figure 3*A***) To implement a model capable of generating double coils that depend upon glutamatergic transmission, we built upon the single coiling model by adding two populations of neurons. We reasoned that if double coiling depended upon excitatory neurotransmission, then a population of commissural excitatory neurons could be necessary to trigger a second contralateral coil in double coils. V0v neurons are a population of glutamatergic commissural interneurons, some of which may be involved in larger amplitude locomotor movements such as coiling (Jay & McLean, 2019). Thus, we added a chain of V0vs (ten neurons for each side) electrically coupled to the previous scaffold (i.e. the ipsilateral IC-MN-V0d scaffold). To generate the crossing excitation underlying the second coil, all V0vs projected glutamatergic synapses to contralateral ICs. Electrical synapses were formed with neighbouring MNs, V0ds, and V0vs (the nearest three of each type of neuron in both the rostral and the caudal directions). Ipsilateral ICs were coupled with V0vs in the first four rostral somites.

**Figure 3.**
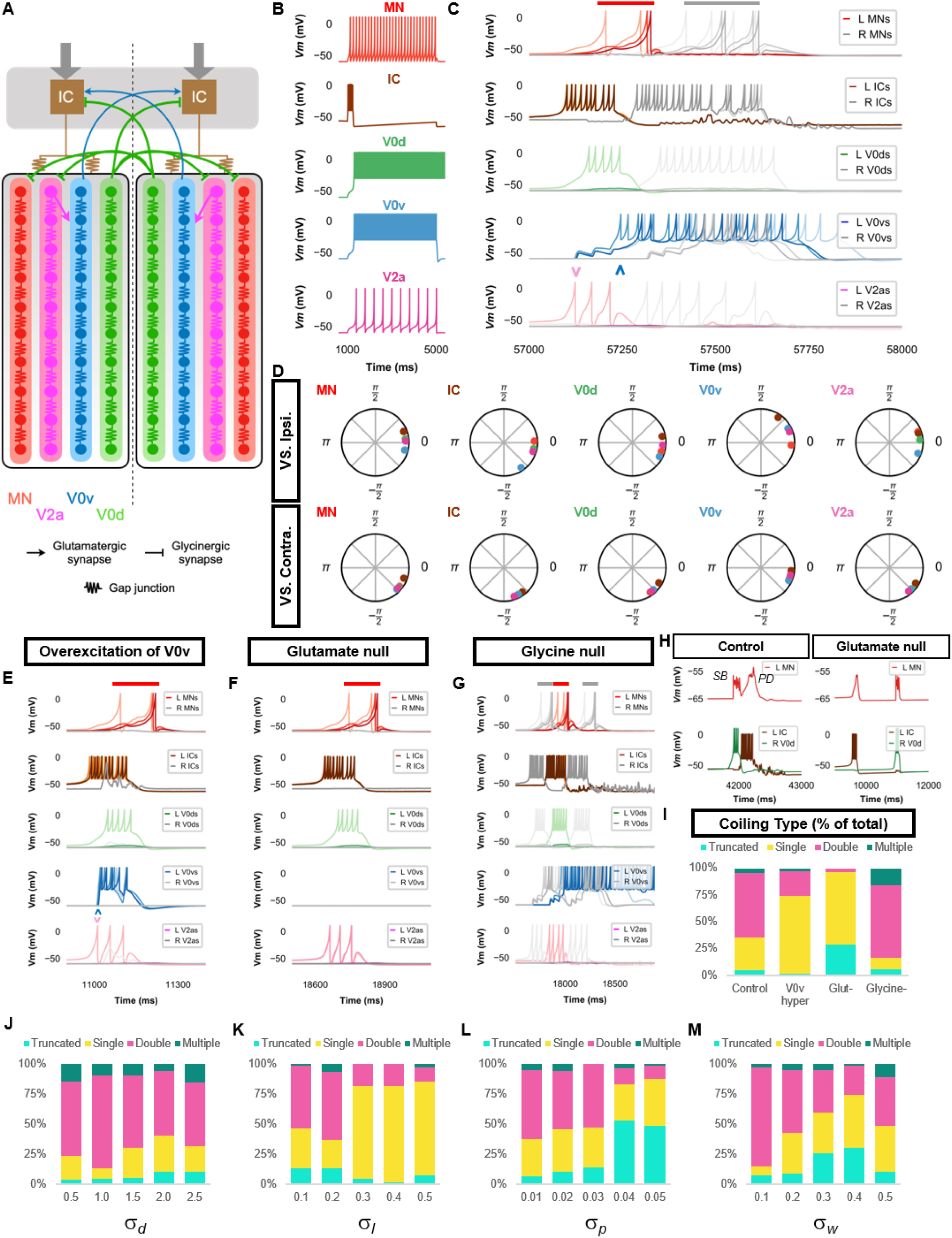
Double coiling model relies on a hybrid network of electrical and chemical synapses. (**A**) Schematic of the double coiling model. Gap junctions between spinal neurons are not depicted. Dashed line indicates the body midline. Gray arrows indicate descending motor command. (**B**) Membrane potential (*Vm*) response of isolated spinal neuron models to a depolarizing current step. (**C**) *Vm* of spinal neurons during a double coil. (**D**) The phase delay of left neurons in relation to ipsilateral and contralateral spinal neurons in the 5^th^ somite and an IC in the rostral kernel during five consecutive left-right double coils. The reference neuron for each polar plot is labelled, and all neurons follow the same color-coding as the rest of the figure. A negative phase delay indicates that the reference neuron precedes the neuron to which it is compared. A phase of 0 indicates that a pair of neurons is in-phase; a phase of π indicates that a pair of neurons is out-of-phase. *Vm* in simulations where (**E**) the weights of the V2a to V0v and the V0v to IC synapses were increased to show that early excitation of V0v prevented the initiation of a second coil following a single coil, (**F**) all glutamatergic transmission was blocked, and (**G**) glycinergic transmission was blocked. (**H)** *Top row*, mixed event composed of a synaptic burst (*SB*) directly followed by a periodic depolarization (*PD*) in a motoneuron in control but not in glutamate null conditions. *Bottom row*, *Vm* in left IC and right V0d during events in top row. (**I**) Proportions of single, double, multiple, and truncated coiling events under control, glutamate null (Glut^-^), overexcited V0vs (V0v hyper), and glycine null (Glycine^-^) conditions. Each condition was tested with five 100,000 ms runs with *σ*_*d*_ = 0.5, *σ*_*p*_ = 0.01, and *σ*_*w*_ = 0.05. (**J**-**M**) Sensitivity testing showing proportions of single, double, multiple and truncated coiling events during ten 100,000 ms runs for each value of *σ*_*d*_, *σ*_*l*_, *σ*_*p*_, and *σ*_*w*_ tested. Solid red and gray bars in **C**,**E-G** indicate the duration of coils. Chevrons in (**C**) and (**E)** denote the initial spiking of V0vs and V2as to indicate latency of V0v firing during the first coil. For **C,E-G**, the *Vm* of a rostral (lightest), middle, and caudal (darkest) neuron is shown, except for IC neurons that are all in a rostral kernel. L: left, R: right. See also ***Figure 3* - *figure supplement 1*** and ***2*** and ***Figure 3* - *video 1-4*.**

The second population of neurons that we added were ipsilaterally projecting excitatory neurons present at this stage and shown to receive mixed PD-SB events (Knogler et al., 2014). These neurons have been suggested to be circumferential ipsilateral descending neurons that arise from the V2a population. In the model, V2as were electrically coupled to IC neurons and projected glutamatergic synapses to V0vs. This chemical synapse caused a delay after the initiation of the initial coil that facilitates the second contralateral coil (see below). V2as most likely also excite motoneurons, based on the data from Knogler et al. (2014). For computational simplicity, we omitted this connection as it was unnecessary for double coilings to be generated, though this may reduce the amplitude of the coils. As V2as display SBs at this stage, we modelled glycinergic projections from V0ds to contralateral V2as such that the *i^th^* V0d projected to all contralateral V2as between the *i-5* and *i+5* segments like how V0ds project to contralateral MNs.

Left and right ICs received a tonic motor command though we delayed the activation of the tonic command to the right side by 1,500 ms to ensure that double coils were not near-coincident bilateral single coils. We modified several parameters of the ICs, most notably increases in the *a* and the *k* parameters, to produce a more extended inter-coiling period (**Figure 3*B***) than seen in single coiling (Knogler et al., 2014). A reminder that the *a* parameter represents the time scale of the recovery variable *u* that returns the membrane potential to rest. The *k* parameter shapes subthreshold dynamics.

SIMULATIONS RESULTS Simulations of the double coiling model frequently generated pairs of successive, left-right alternating coils lasting about one second in total (**Figure 3*C***, **Figure 3 - video 1**). In five 100,000-ms long runs of the base model with a minimal amount of variability added to several model parameters (*σ*_*d*_ = 0.5, *σ*_*p*_ = 0.01, and *σ*_*w*_ = 0.05), approximately 60% of events were double coils, with the rest mainly being single coils (31%), and very few triple coils or truncated single coils (5% each) (**Figure 3*I***).

The timing of ICs, V2as, and V0vs (**Figure 3*C***) suggest that double coils were generated by ipsilateral recruitment of V2as and V0vs during the first coil, which led to activation of the contralateral ICs to initiate the second coil. This sequence is supported by an analysis of the phase delays (**Figure 3*D***). IC firing precedes the firing of all other ipsilateral spinal neurons suggesting they drive the activity of each coil. V2a activity precedes that of V0vs, which suggests that V2as recruit V0vs. This recruitment of V0vs by V2as is supported by simulations where the V2a to V0v synapse is removed (**Figure 3 - figure supplement 1*A***). V0v activity succeeds all other ipsilateral spinal interneurons, suggesting they are the last interneurons active during the first coil in a double coiling event. A key to generating double coils in our model was thus to delay the activation of V0vs. This delay enabled the activation of contralateral ICs after the first coiling is completed and when commissural inhibition of the contralateral IC has also terminated. If the activation of ipsilateral V0vs occurred too early during the first coiling, which can be produced by increasing the weight of the V2a to V0v and the V0v to contralateral IC glutamatergic synapses, the occurrence of a second coil is less likely (**Figure 3*E,I***, **Figure 3 - video 2**).

To further underscore the importance of glutamatergic transmission to double coiling as reported experimentally (Knogler et al., 2014), blocking glutamatergic transmission in the model greatly reduced the number of double coils (**Figure 3*F,I***, **Figure 3 - video 3**). On the other hand, blocking glycinergic synapses increased multiple coilings of three or more coils (**Figure 3*G,I***, **Figure 3 - video 4**) as Knogler et al. (2014) reported. This effect presumably occurs due to the unopposed reverberating commissural excitation of ICs by V0vs. Indeed, silencing V0v synapses in a model with no glycinergic synapses blocks double and multiple coils (**Figure 3 - figure supplement 1*B***).

The sequencing of commissural excitation and inhibition in the generation of double coils is further underscored by the presence of mixed SB-PD or PD-SB events (**Figure 3*H***) observed experimentally in hyperpolarized MNs (Knogler et al. 2014). In these mixed events, the PDs were generated by gap junction coupled ICs during the ipsilateral coil, whereas contralateral V0ds activated during the contralateral coil generated the SBs in the ipsilateral MNs. Blocking glutamatergic transmission in our model uncoupled PDs and SBs (**Figure 3*H***) as observed experimentally (Knogler et al. 2014).

Just as the robustness of the single coiling model was tested through modifications to the base model and several sensitivity tests, we also tested the robustness of the double coiling model. First, we increased the size of the model from ten to thirty somites. Modifications of the tonic motor command amplitude, length of IC axons, gap junction coupling from IC to MN, and the synapses from MN to muscle cell, V0v to IC, V2a to V0v enabled the generation of double coils in this model (**Figure 3 - figure supplement 1*C*, Figure 3 - video 5**). In the ten-somite base model, we also tested the role of the glycinergic reversal potential. The value of this parameter was hyperpolarized from -45 mV in the single coiling model to -58 mV in the base double coiling model. This shift was intended to reflect gradual hyperpolarization of the reversal potential of glycine during development (Ben-Ari, 2002, p.; Saint-Amant & Drapeau, 2000, 2001). We tested the double coiling model at values ranging between -46 to -70 mV (**Figure 3 - figure supplement 2*A***). We found that the proportion of double coils seemed to be higher, and the proportion of multiple coils was increased at more depolarized values of the glycinergic reversal potential. The proportion of double coils was relatively constant at more hyperpolarized values of the glycinergic reversal potential.

To test whether the double coiling model was sensitive to within-model parameter variability, we ran sets of ten 100-s long simulations at various values of *σ*_*d*_, *σ*_*l*_, *σ*_*p*_, and *σ*_*w*_(**Figure 3*J*-*M***). Again, we found that relatively small levels of variability in the parameters governing membrane dynamics (*σ*_*p*_) decreased the proportion of coiling events that were double coils and increased the number of truncated coils (**Figure 3*L***). Moderate levels of variability in the parameters governing axonal length (*σ*_*l*_) or synaptic weight (*σ*_*w*_) decreased the proportion of double coils while increasing single coils and sometimes truncated coils (**Figure 3*K*,*M***). The proportion of coiling events was largely unaffected by variability in the amplitude of the motor command (*σ*_*d*_, **Figure 3*J***).

Considering that the generation of double coils was sensitive to chemical synaptic activity and gap junctions (Knogler et al. 2014), we tested the sensitivity of the model to variability in the weights of only chemical synapses (*σ*_*w,chem*_) and only gap junctions (*σ*_*w,gap*_). We found that the proportion of double coils was relatively more sensitive to the variability of gap junctions than chemical synapses (**Figure 3 - figure supplement 2*B,C***).

### Generation of swimming pattern by spinal network oscillators

Around 2 or 3 dpf, zebrafish transition from coiling movements to swimming (Drapeau et al., 2002; Saint-Amant & Drapeau, 1998). This transition entails two fundamental changes in locomotor movements: First, long, slow coils are replaced by quick, short tail beats; and secondly, the number of consecutive tail beats are increased from the two side-to-side coilings seen in double coils to multiple consecutive tail beats that compose each swimming episode. One of the emerging swimming movements is beat-and-glide swimming, characterized by short swimming episodes lasting several hundreds of ms separated by gliding pauses and lasting several hundreds of ms (Budick & O’Malley, 2000; Buss & Drapeau, 2001). Swim episodes consist of repetitive left-right alternating, low-amplitude tail beats that propagate from the rostral toward the caudal end of the fish body and are generated approximately at 20 to 65 Hz (Budick & O’Malley, 2000; Buss & Drapeau, 2001).

Beat-and-glide swimming can be produced in isolated larval zebrafish spinal cord preparations by NMDA application (Lambert et al., 2012; McDearmid & Drapeau, 2006; Wiggin et al., 2012) or by optogenetic stimulation of excitatory spinal neurons (Wahlstrom-Helgren et al., 2019). This capacity suggests that the transition from coiling to swimming involves a delegation of rhythm generation from ICs to spinal locomotor circuits (Roussel et al., 2020). Therefore, we sought to model a spinal network that could generate beat-and-glide swimming activity hallmarks - swim episodes lasting about 200-300 ms with repeated left-right alternating low-amplitude tail beats at around 20-65 Hz – without relying on pacemaker cells.

Recent experimental studies have also started to delineate the contributions of specific populations of spinal neurons to swimming. Ablation of ipsilaterally projecting, excitatory neurons in the V2a population eliminates swimming activity (Eklof-Ljunggren et al., 2012). Genetic ablation of ipsilaterally projecting, inhibitory neurons in the V1 population affects swim vigor but has no effects on the patterning of swimming (Kimura & Higashijima, 2019). Genetic ablation of a subset of commissural inhibitory neurons in the dI6 population reduces left-right alternation (Satou et al., 2020). We sought to replicate the role of these neurons in our model.

NETWORK DESCRIPTION (**Figure 4*A***) Whereas coiling is likely to be generated by primary motoneurons, swimming is more likely to involve secondary motoneurons (Ampatzis et al., 2013; D. W. Liu & Westerfield, 1988). There are more secondary than primary motoneurons, and new spinal neurons are born at the same time as secondary motoneurons (Bernhardt et al., 1990). To emulate the increase in the number of spinal neurons that may underlie swimming, we increased the size of the fish from ten to fifteen segments and accordingly increased the number of MNs, V0vs, and V2as from ten to fifteen. Thus, each model somite in our swimming model represented two biological somites instead of three in our coiling models. IC neurons were removed from the model to reduce computational load. We are not aware of any experimental evidence of the involvement of IC neurons in later swimming stages.

**Figure 4.**
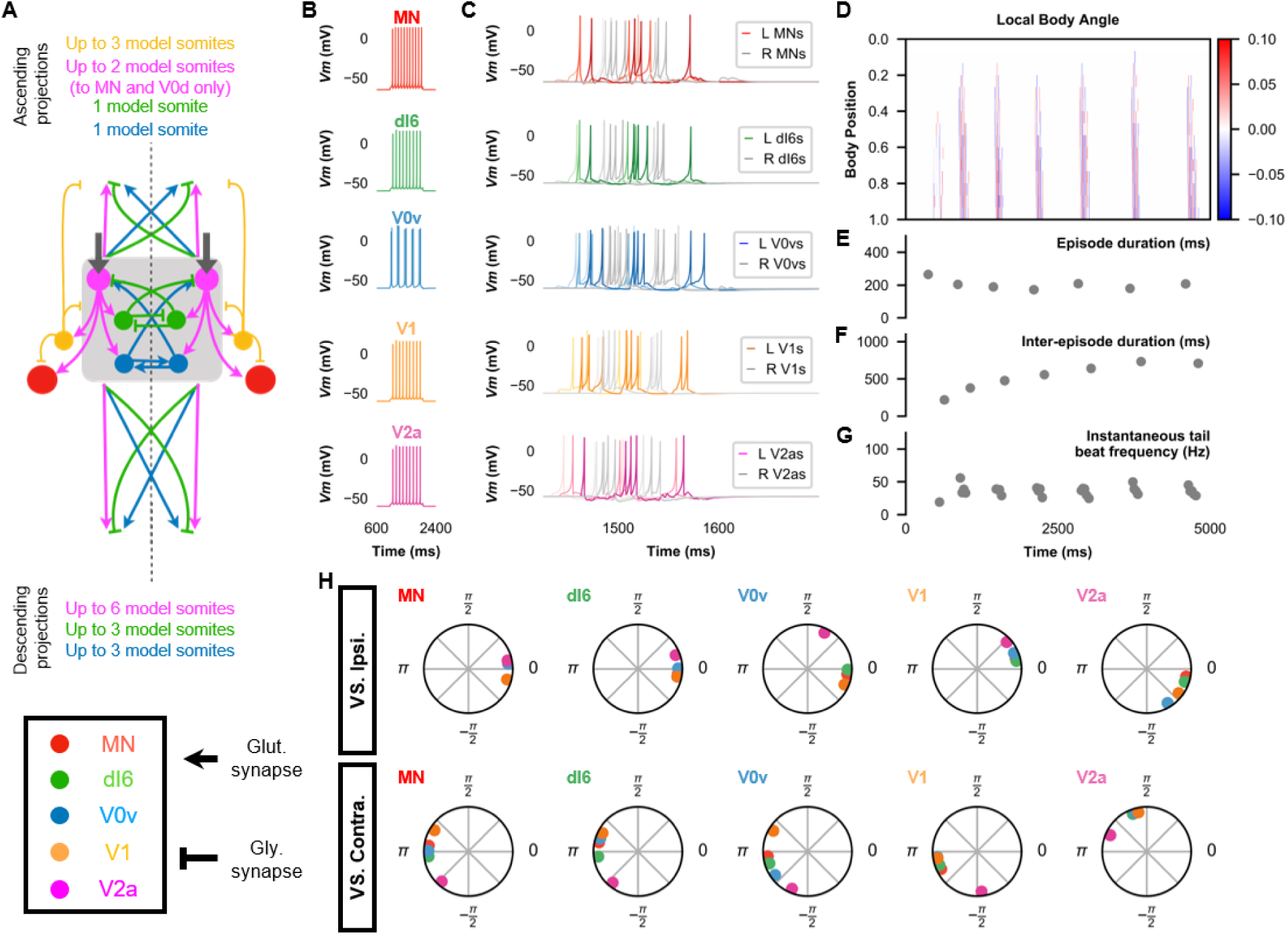
The base model for beat-and-glide swimming. (**A**) Schematic of the model architecture underlying beat-and-glide swimming. (**B**) Membrane potential (*Vm*) response to a depolarizing current step of isolated spinal neurons in the model. (**C**) *Vm* of spinal neurons during a beat-and-glide swimming simulation. The *Vm* of a rostral (lightest), middle, and caudal (darkest) neuron is shown. L: left, R: right. (**D)** Heat-map of local body angle. (**E**) Episode duration, (**F**) inter-episode interval, (**G**) instantaneous tail beat frequency, and (**H**) the phase delay of left neurons in relation to ipsilateral and contralateral spinal neurons in the 10^th^ somite during a 10,000 ms simulation. The reference neuron for each polar plot is labelled, and all neurons follow the same color-coding as the rest of the figure. A negative phase delay indicates that the reference neuron precedes the neuron to which it is compared. A phase of 0 indicates that a pair of neurons is in-phase; a phase of π indicates that a pair of neurons is out-of-phase. See also ***Figure 4* - *video 1*.**

We introduced two populations of neurons for the beat-and-glide model. Commissural inhibitory CoBL neurons (including neurons from the dI6 and V0d populations) are active during swimming (Liao & Fetcho, 2008; Satou et al., 2020). V0d neurons were involved with faster swimming, whereas dI6s were more likely to be active during slower swimming (Satou et al., 2020). Therefore, we modelled CoBL neurons as dI6s. The dI6s thus replaced the V0ds in the coiling models as the source of contralateral inhibition in the swimming model. CoBL neurons have been shown to project to motoneurons, dI6s, and unidentified ipsilateral descending spinal neurons that could be V2as (Satou et al., 2020). We added fifteen dI6s per side. We modelled the projection pattern of dI6s based on the bifurcating axons with short ascending and long descending branches of a subset of neurons in the dI6 subpopulation (Satou et al., 2020). Thus, the *i*^th^ dI6s projected ascending branches to their rostral targets in the *i-1*th segment and projected descending branches to their caudal targets between the *i+1* and *i+3* segments.

A second new population of neurons was the V1 interneurons that include circumferential ascending (CiA) interneurons that emerge during the second wave of neurogenesis in the spinal cord (Bernhardt et al., 1990). While V1 neurons were not included in the coiling models because their role in that form of movement is unclear, experiments in which the genetically identified V1 neurons are ablated suggest a role in controlling swim vigor (Kimura & Higashijima, 2019). We thus modelled V1s as a population of tonic firing neurons with ipsilateral ascending glycinergic projections. We added fifteen V1s per side. We distributed V1s from segment 2 to the caudal end because our preliminary simulations suggested that starting the distribution of V1s at segment 1 made the episode duration more variable. In our model, V1s project segmental and ascending ipsilateral glycinergic synapses with rostral V2as (Kimura & Higashijima, 2019), dI6s, and V0vs (Sengupta et al., 2021) such that the V1s in the *i*^th^ segment project to their rostral targets in the *i-1* to *i-2* segments. The V1 projections were short based upon recent evidence that their projections to motor circuits are constrained to segmental and immediately-neighbouring somites (Sengupta et al., 2021). Reciprocally, V2as formed glutamatergic synapses to caudally located V1.

V2as were considered the primary source of rhythmogenesis in our models of beat-and-glide swimming based on previous studies showing the necessity and sufficiency of V2a neurons to swimming activity (Eklof-Ljunggren et al., 2012; Ljunggren et al., 2014). In the swimming model, V2as projected segmental and descending projections to dI6s, MNs, V0vs, V1s, and caudal V2as (the *i^th^* V2a projected to all caudal V0vs, dI6s, V2as, and MNs between the *i+1* and *i+6* segments). Connections between V2as and V2as (Ampatzis et al., 2014; Menelaou & McLean, 2019; Song et al., 2020) and from V2a neurons to MNs (Ampatzis et al., 2014; Menelaou & McLean, 2019; Song et al., 2020) have been reported. Connections from V2as to dI6s have not been studied yet, and so we based them on the reported connections from V2a neurons to V0d neurons, another population of commissural inhibitory interneurons (Menelaou & McLean, 2019). A subtype of V2a neurons that project to MNs was shown to bifurcate and have short ascending branches (Menelaou & McLean, 2019), which we modelled in addition to an ascending V2a to V0v connection (the *i^th^* V2a projected to rostral V0vs and MNs in the *i-1* and *i-2* segments) that remains to be confirmed.

Less is known about the connection patterns of commissural excitatory neurons at larval stages. However, in adult zebrafish, V0v commissural excitatory neurons have been shown to have bifurcating axons with shorter ascending branches and longer descending branches (Björnfors & El Manira, 2016). We modelled V0vs to project only to contralateral V2as. The *i*^th^ V0vs projected ascending branches to their rostral targets in the *i-1*^th^ segment and projected descending branches to their caudal targets between the *i+1* and *i+3* segments.

Whether rhythmic motor commands from supraspinal commands generate rhythmic tail beats at the spinal cord level is unclear. There is evidence for rhythmic and tonic activity in reticulospinal neurons involved in swimming (Kimura et al., 2013). However, the isolated zebrafish spinal cord can generate rhythmic activity similar to swimming (McDearmid & Drapeau, 2006; Wahlstrom-Helgren et al., 2019; Wiggin et al., 2012, 2014). Therefore, in our model, V2as received a tonic motor command in the form of a DC current to test whether the rhythm and pattern of swimming could be generated solely from the activity of spinal circuits.

Most spinal neurons at larval stages exhibit either tonic firing or firing with spike rate adaptation (Kimura & Higashijima, 2019; Menelaou & McLean, 2012, 2019; Satou et al., 2020), while a subset of motoneurons showing intrinsic burst firing (Menelaou & McLean, 2012). Therefore, we posited that the generation of rhythmic tail beats was not dependent upon the presence of intrinsically bursting neurons. Almost all of the neurons in the beat-and-glide swimming model were modelled to fire tonically (**Figure 4*B***). Our base model was able to generate the beat-and-glide swimming pattern - alternating episodes of tail beats followed by silent inter-episode intervals each lasting hundreds of seconds - if V0vs were modelled to exhibit a more chattering or bursting firing pattern (**Figure 4*B***).

As the model is symmetrical, including the motor command, we found that the model produces synchronous left-right activity unless we introduced some variability in commissural connections. With no *a priori* knowledge of where such variability could arise from, we chose to introduce a small amount of variability in the contralateral inhibition of dI6s by dI6s. The synaptic weights of this connection were scaled by a random number picked from a Gaussian distribution with mean, µ = 1, and standard deviation of 0.1, and this was sufficient to generate alternating left-right alternation. We did not seek to further characterize the variability required to generate left-right alternation.

SIMULATION RESULTS The beat-and-glide swimming model exhibited short-duration (hundreds of ms) swimming episodes with left-right alternation and tail beat frequencies between 20-60 Hz (**Figure 4*C-G*, Figure 4 - video 1**). The characteristics of the swimming episodes in our simulations were close to those described for free swimming in larval zebrafish by Buss and Drapeau (2001), though the swimming output in our simulations had larger episode durations, shorter inter-episode intervals, and lower tail beat frequencies (**Table 1**).

**Table 1.** Comparison of beat-and-glide swimming in model and experimental data from Buss and Drapeau (2001)

| *Parameter* | *Base model* | *Buss and Drapeau (2001)* |
| --- | --- | --- |
| *Mean episode duration (ms)* | 234 +/- 6 | 180 +/- 20 |
| *Mean inter-episode interval (ms)* | 242 +/- 20 | 390 +/- 30 |
| *Mean tail beat frequency (Hz)* | 30.0 +/- 0.6 | 35 +/- 2 |

Values in mean +/- standard error. $n = \text{ten } 10,000 \text{ ms-long simulations for the base model, and } n = 12 \text{ animals for the data from Buss and Drapeau (2001).}$

An analysis of the phase delay between neuron populations during beat-and-glide swimming in the base model shows that the activity of ipsilateral, glutamatergic V2as precedes the activity of all ipsilateral neurons (**Figure 4*H***). This earlier firing of V2as suggests that these spinal neurons drive the activity of the ipsilateral spinal swimming circuit. On the other hand, the glycinergic V1 neurons succeed all ipsilateral spinal neurons, suggesting they provide negative feedback to ipsilateral spinal swimming circuits. The contralateral dI6s and V0vs are out-of-phase with contralateral spinal neurons, consistent with their role in mediating left-right coordination. The longest delay between V2as and their ipsilateral counterparts was with the V0vs, which is reminiscent of V2a firing preceding V0v firing in the double coiling model to ensure a sufficient delay for the initiation of the second coil. Thus, the generation of alternating left-right tail beats would seem to require a certain delay in the excitation of contralateral swimming circuits.

To further investigate the role of specific neurons in the model’s swimming activity, we performed simulations composed of three 5,000 ms long epochs: Epoch 1, where the model was intact; Epoch 2, where we silenced the targeted neurons by removing their synaptic inputs; and Epoch 3, where the synaptic inputs to the targeted neurons were restored. Silencing V2as abolished the generation of tail beats (**Figure 5*A-F***, **Figure 5 - figure supplement 1*A-F*)**, underscoring their primacy to the generation of tail beats (Eklof-Ljunggren et al., 2012; Ljunggren et al., 2014). Commissural excitation mediated by V0vs seemed to be very important in maintaining the beat-and-glide pattern. Silencing V0vs diminished but did not eliminate the rhythmic firing of V2as or MNs. During Epoch 2, the tonic motor command continued to activate V2as. Pairs of left-right tail beats may result from the commissural inhibition by dI6s that was still present. However, removing the contralateral excitation by V0v prevented the repetitive activation of the silent side after each tail beat, which severely reduced episode duration and the number of tail beats generated in each episode (**Figure 5*G-L***, **Figure 5 - figure supplement 1*G-L***, **Figure 5 - video 1**). These simulations suggest that commissural excitation is necessary to repeatedly activate the silent contralateral side during ongoing swimming to ensure successive left-right alternating tail beats and longer swim episodes (Björnfors & El Manira, 2016; Saint-Amant, 2010).

**Figure 5.**
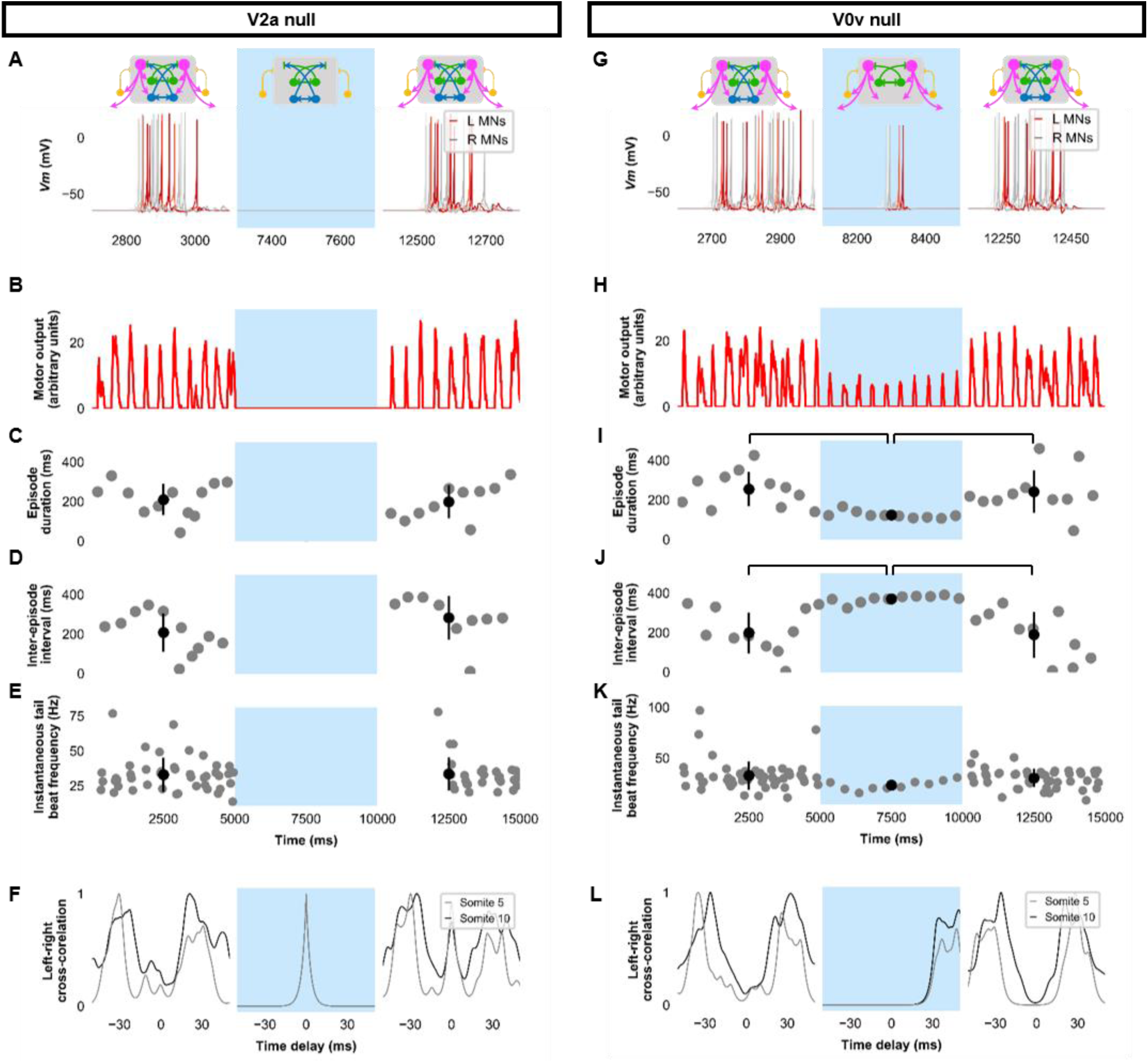
Silencing spinal excitatory neurons during beat-and-glide swimming. Simulations consisted of three 5,000 ms epochs. In the middle epoch, silencing of targeted spinal neurons was achieved by removing all synaptic and external currents from the targeted population. Synaptic and external currents were restored in the last epoch. (**A*-*F**) Simulations where V2as were silenced and (**G-L**), where V0vs were silenced. (**A**, **G**) *Top*, the functional state of the spinal network during the three epochs. *Bottom*, Motoneuron (MN) membrane potential (*Vm*) during simulations where targeted neurons were silenced in the middle epoch. The *Vm* of a rostral (lightest), middle, and caudal (darkest) neuron is shown. (**B**, **H**) The integrated muscle output, (**C**, **I**) episode duration, (**D**, **J**) inter-episode intervals, and (**E**, **K**) instantaneous tail beat frequency during each respective simulation. Averages within epoch are shown in black (mean ± s.d.). Brackets denote significant pairwise differences. (**F**, **L**) the left-right coordination of somites 5 and 10. L: left, R: right. The first part of epoch 3 of the V2a silenced simulation involved synchronous left-right activity, hence the lack of instantaneous tail beat frequency values. ***Statistics***: For (**C**-**E**), there were no episodes during epoch 2. There were no statistically significant differences between epoch 1 and 3 for any of the parameters. (**I**) *F*_2,31_ = 7.2, p = 0.0029. (**J**) *F*_2,28_ = 10.2, p = 0.001. (**K**) *F*_2,115_ = 3.0, p = 0.055. P-values for t-tests are found in ***Figure 5* – *source data 1***. See also ***Figure 5* - *figure supplement 1*** and ***Figure 5* - *video 1*** and ***2*.**

Genetic ablation of the ipsilaterally ascending inhibitory V1 interneurons increased swim vigor but produced no overt changes in swimming patterns (Kimura and Higashijima 2019). Consistent with those experimental results, simulating the removal of ipsilateral ascending inhibition by silencing V1 interneurons seemed to increase the amplitude of motoneuron activity (**Figure 6*A,B***, **Figure 6 - figure supplement 1*A-F***, **Figure 6 - video 1**). While the overall beat-and-glide pattern persisted, the duration of episodes was increased, and inter-episode intervals were shortened between Epochs 1 and 2 (**Figure 6*C*,*D***). Tail beat frequency was increased (**Figure 6*E***). Left-right alternation was reduced in these simulations and during simulations where dI6s were silenced (**Figure 6*F,G,L***, **Figure 6 - figure supplement 1*G-L***). The reduction in left-right alternation during simulations where dI6s were silenced was greater in caudal somites than in rostral somites (**Figure 6*L***). Note that while left-right alternation was reduced, this did not preclude left-right alternating tail beats from being generated (**Figure 6 - video 2**). The reduction of left-right coordination seen here was comparable to levels seen after genetic ablation of a commissural inhibitory subpopulation of dI6 interneurons (Satou et al., 2020) but is not sufficient to prevent left-right alternation. Since swimming is generated by rostrocaudal propagation of contractile waves, any left-right alternation in rostral segments will inevitably sway the rest of the body, as suggested by the musculoskeletal model. The precise kinematics of the tail beats will be affected by the reduction of left-right alternation observed (**Figure 6 - figure supplement 2, Figure 6 - video 2**). Finally, silencing dI6s had negligible effects on the episode duration and inter-episode interval but increased tail beat frequency when comparing Epochs 1 and 2 and may increase the amplitude of motor activity (**Figure 6*H*-*K***).

**Figure 6.**
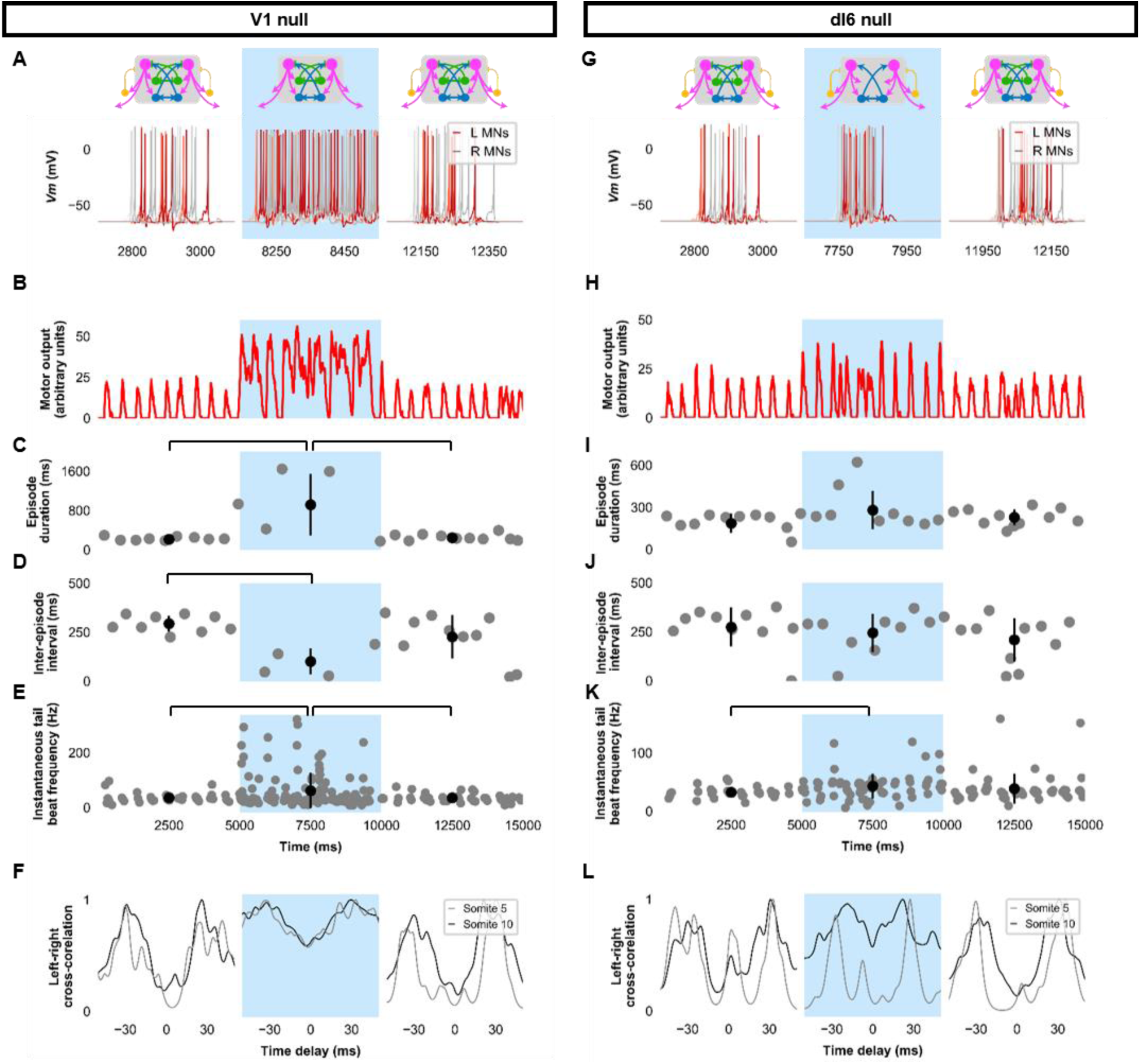
Silencing spinal inhibitory neurons during beat-and-glide swimming. Simulations consisted of three 5,000 ms epochs. In the middle epoch, silencing of targeted spinal neurons was achieved by removing all synaptic and external currents from the targeted population. Synaptic and external currents were restored in the last epoch. (**A*-*F**) Simulations where V1s were silenced and (**G-L**), where dI6s were silenced. (**A**, **G**) *Top*, the functional state of the spinal network during the three epochs. *Bottom*, Motoneuron (MN) membrane potential (*Vm*) during simulations where targeted neurons were silenced in the middle epoch. The *Vm* of a rostral (lightest), middle, and caudal (darkest) neuron is shown. (**B**, **H**) The integrated muscle output, (**C**, **I**) episode duration, (**D**, **J**) inter-episode intervals, and (**E**, **K**) instantaneous tail beat frequency during each respective simulation. Averages within epoch are shown in black (mean ± s.d.). Brackets denote significant pairwise differences. (**F**, **L**) the left-right coordination of somites 5 and 10. L: left, R: right. ***Statistics***: (**C**) *F*_2,25_ = 10.5, p = 5.8 x 10^-4^. (**D)** *F*_2,22_ = 6.6, p = 0.0063. (**E**) *F*_2,214_ = 6.9, p = 0.0013. (**I**) *F*_2,31_ = 2.5 p = 0.10. (**J**) *F*_2,28_ = 0.9, p = 0.42. (**K**) *F*_2,145_ = 3.5, p = 0.033. P-values for t-tests are found in ***Figure 6* – *source data 1***. See also ***Figure 6* - *figure supplement 1*** and **2** and ***Figure 6* - *video 1*** and ***2*.**

Previous experimental results showed that strychnine application disturbed swimming in the 20- 40 Hz range at later stages of development (Roussel et al., 2020). Our model’s behaviour to loss of glycinergic transmission was tested. Removal of all glycinergic transmission led to continual tail beats with minimal gliding periods (**Figure 7**). Motor output was increased during the epoch of no glycinergic transmission (**Figure 7*B***), episode duration was considerably lengthened, and the inter-episode interval was shortened (**Figure 7*C*-*D***). The frequency of tail beats increased (**Figure 7*E***). Left-right alternation was reduced, particularly at caudal somites (**Figure 7*F***), which did not preclude left-right tail beats but altered swimming kinematics (**Figure 7 – figure supplement 1**, **Figure 7 - video 1**). These results indicate that removing glycinergic transmission in the model led to near-continuous swimming activity with altered kinematics and greater frequencies of tail beats.

**Figure 7.**
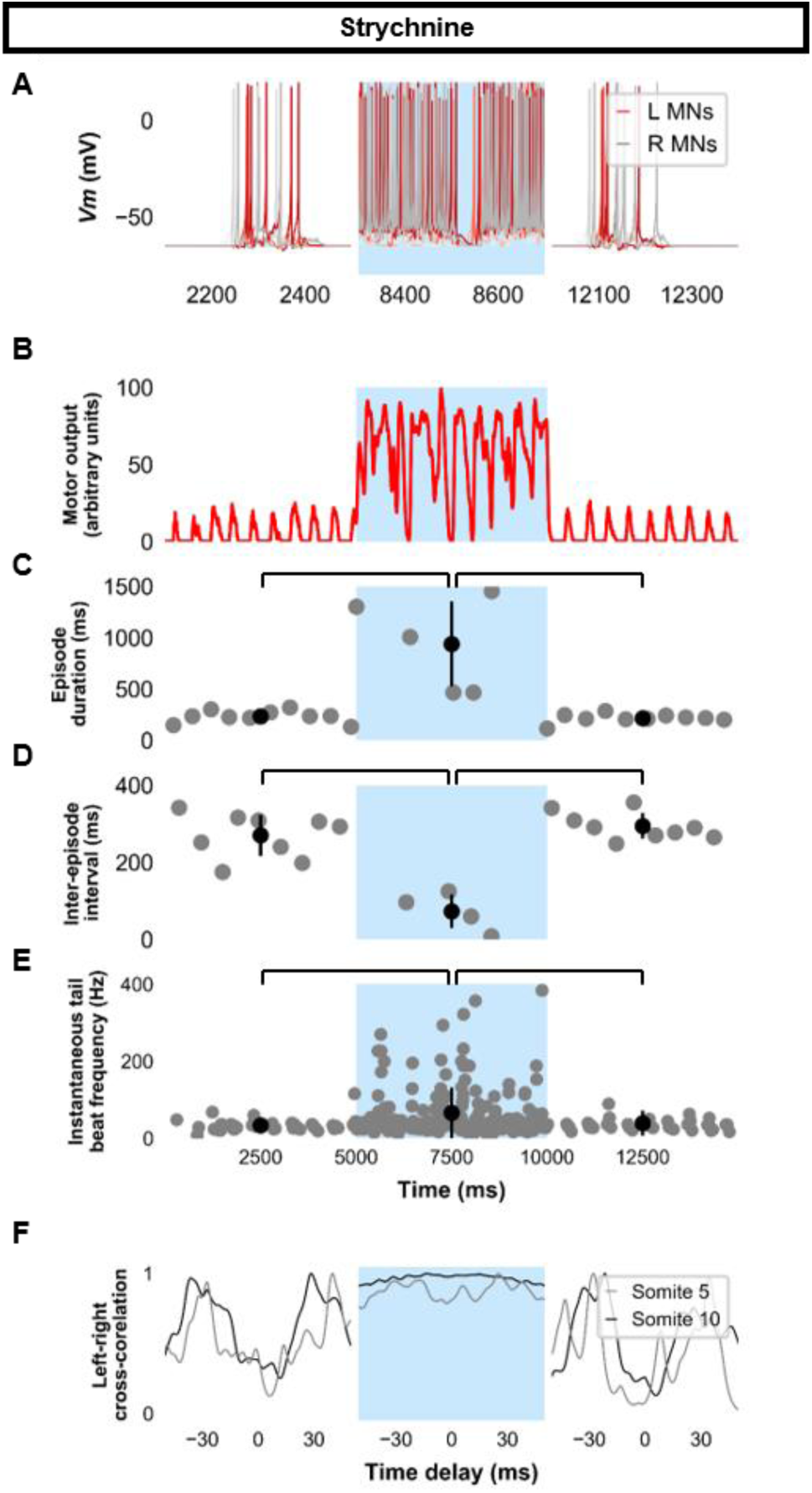
Simulating the effects of strychnine on beat-and-glide swimming. Simulations to assess the effects of blocking glycinergic transmission consisted of three 5,000 ms epochs. In the middle epoch, all glycinergic currents were blocked. Glycinergic transmission was restored in the last epoch. (**A**) Motoneuron (MN) membrane potential (*Vm*) during simulations where glycinergic transmission was blocked in the middle epoch. The *Vm* of a rostral (lightest), middle, and caudal (darkest) neuron is shown. (**B**) The integrated muscle output, (**C**) episode duration, (**D**) inter- episode intervals, and (**E**) instantaneous tail beat frequency during this simulation. Averages within epoch are shown in black (mean ± s.d.). (**F**) The left-right coordination of somites 5 and 10. L: left, R: right. ***Statistics***: (**C**) *F*_2,24_ = 2.5, p = 2.2 x 10^-6^. (**D**) *F*_2,21_ = 32.0, p = 8.3 x 10^-7^. (**E**) *F*_2,267_ = 8.3, p = 0.0003. P-values for t-tests are found in ***Figure 7* – *source data 1*.** See also See also ***Figure 7* - *figure supplement 1* and *Figure 7* - *video 1***

We then proceeded to test the sensitivity of the base model to several model parameters. Since some V2a interneurons in adult zebrafish have pacemaker capacities (Song et al., 2020), we tested whether we could also generate beat-and-glide swimming in a model with bursting V2a (**Figure 8*A***). A model with bursting V2as where we also decreased the strength of synapses from V2as to other neurons and increased the connection strength of V0vs and dI6s to contralateral V2as (**Table 4**) generated 100-400 ms swimming episodes of left-right alternating tail beats at frequencies around 20-80 Hz interspersed by 100-400 ms long silent inter-episode intervals (**Figure 8*B-F*, Figure 8 - video 1**). Surprisingly, a model with only tonic firing neurons (**Figure 8*G***) was able to generate the hallmarks of beat-and-glide swimming as well (**Figure 8*H-L*, Figure 8 - video 2**). While there were eventually longer episodes with shorter inter-episode intervals after 6,000 ms, this simulation suggests that the architecture of the network is sufficient to generate beat-and-glide swimming for long periods despite the absence of any neurons with bursting properties.

**Figure 8.**
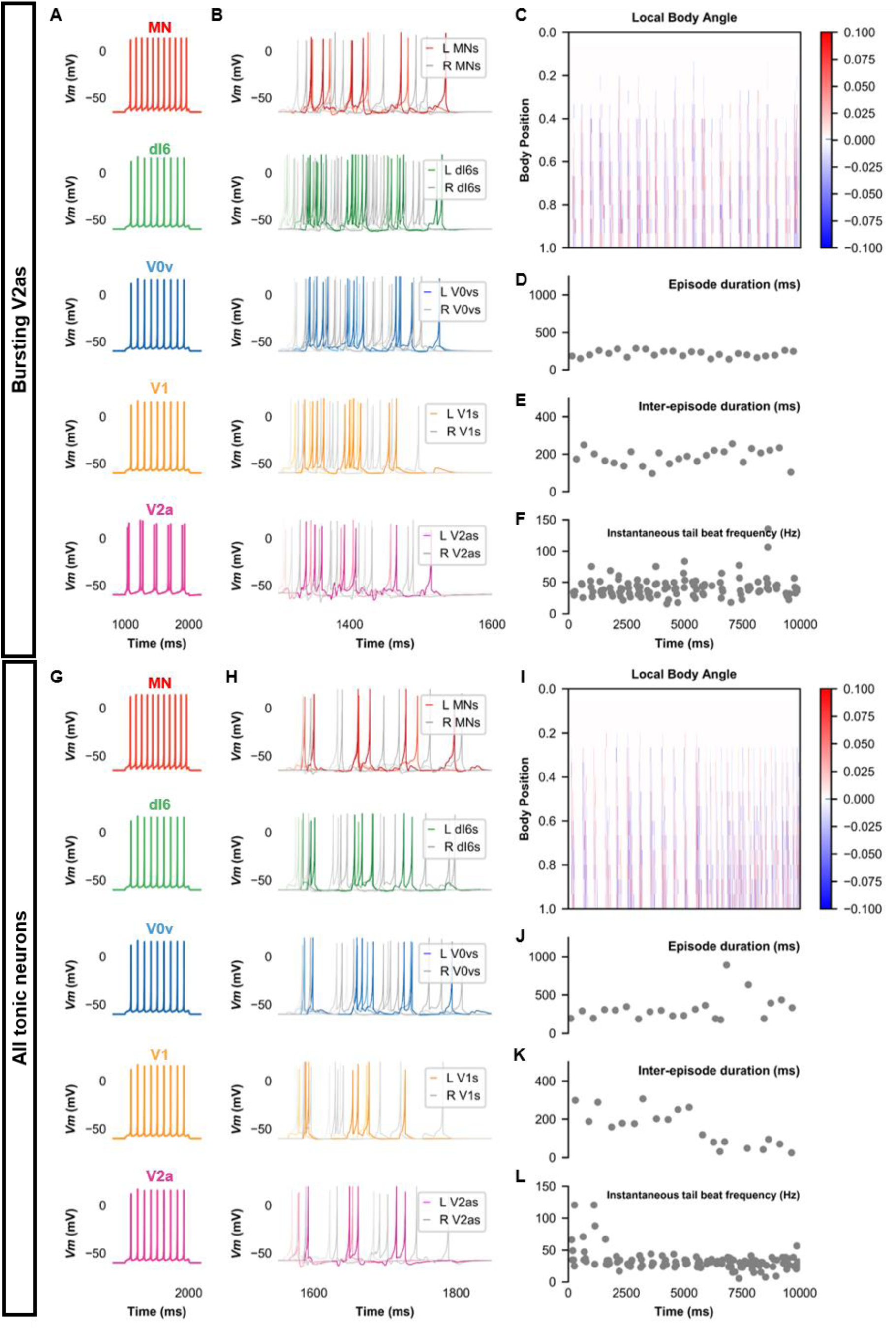
Beat-and-glide models with bursting V2a (A-F) or only tonic neurons (G-L). (**A, G**) Membrane potential (*Vm*) response of isolated neurons in the model to a current step. (**B, H**) *Vm* of spinal neurons during swimming simulation. The membrane potential of a rostral (lightest), middle, and caudal (darkest) neuron is shown. L: left, R: right. (**C, I**) Heat-map of local body angle. (**D, J**) Episode duration, (**E, K**) inter-episode interval, and (**F, L**) instantaneous tail beat frequency during the same simulations as **B** and **H**, respectively. See also ***Figure 8* - *figure supplement 1*** and **2,** and ***video 1*** and ***2***.

We could generate beat-and-glide swimming in a thirty-somite and a ten-somite model (**Figure 8 figure supplement 1, Figure 8 - video 3**). We also examined the effects of changing the glycinergic reversal potential that was set at -70 mV for the base beat-and-glide model (**Figure 8 figure supplement 2**). The glycinergic reversal potential was set at values between -72 mV and -56 mV (**Figure 8 - figure supplement 2*A,B***). Episode duration was increased, and inter-episode intervals decreased at more depolarized values of the glycinergic reversal potential leading to the loss of the beat-and-glide pattern (**Figure 8 - figure supplement 2*C***). Tail beat frequency also increased as glycinergic currents decreased at depolarized glycinergic reversal potentials, and left-right alternation was replaced by left-right synchrony (**Figure 8 - figure supplement 2*D***).

Finally, the sensitivity of the base model to variability was tested by running sets of ten 10,000- ms long simulations at various values of *σ*_*d*_, *σ*_*l*_, *σ*_*p*_, and *σ*_*w*_. We also performed ten 10,000-ms long simulations of the base model (a reminder that there is a random scaling factor to the dI6 to contralateral dI6 synapse in the base model). The episode duration, inter-episode interval, average tail beat frequency in each episode, and the minimum coefficient of the cross-correlation of the left and right muscle output were analyzed (**Figures 9 and 10**). Increases in variability in the motor command drive (*σ*_*d*_) seemed to affect inter-episode intervals and left-right alternation but not episode duration and tail beat frequency (**Figure 9*A*-*D***). At similar levels of variability in rostrocaudal axonal length (*σ*_*l*_), all four measures of swimming activity were perturbed (**Figure 9*E*-*H***). On the other hand, variability in synaptic weights (*σ*_*w*_) affected only episode duration and inter-episode duration (**Figure 9*I*-*L***).

**Figure 9.**
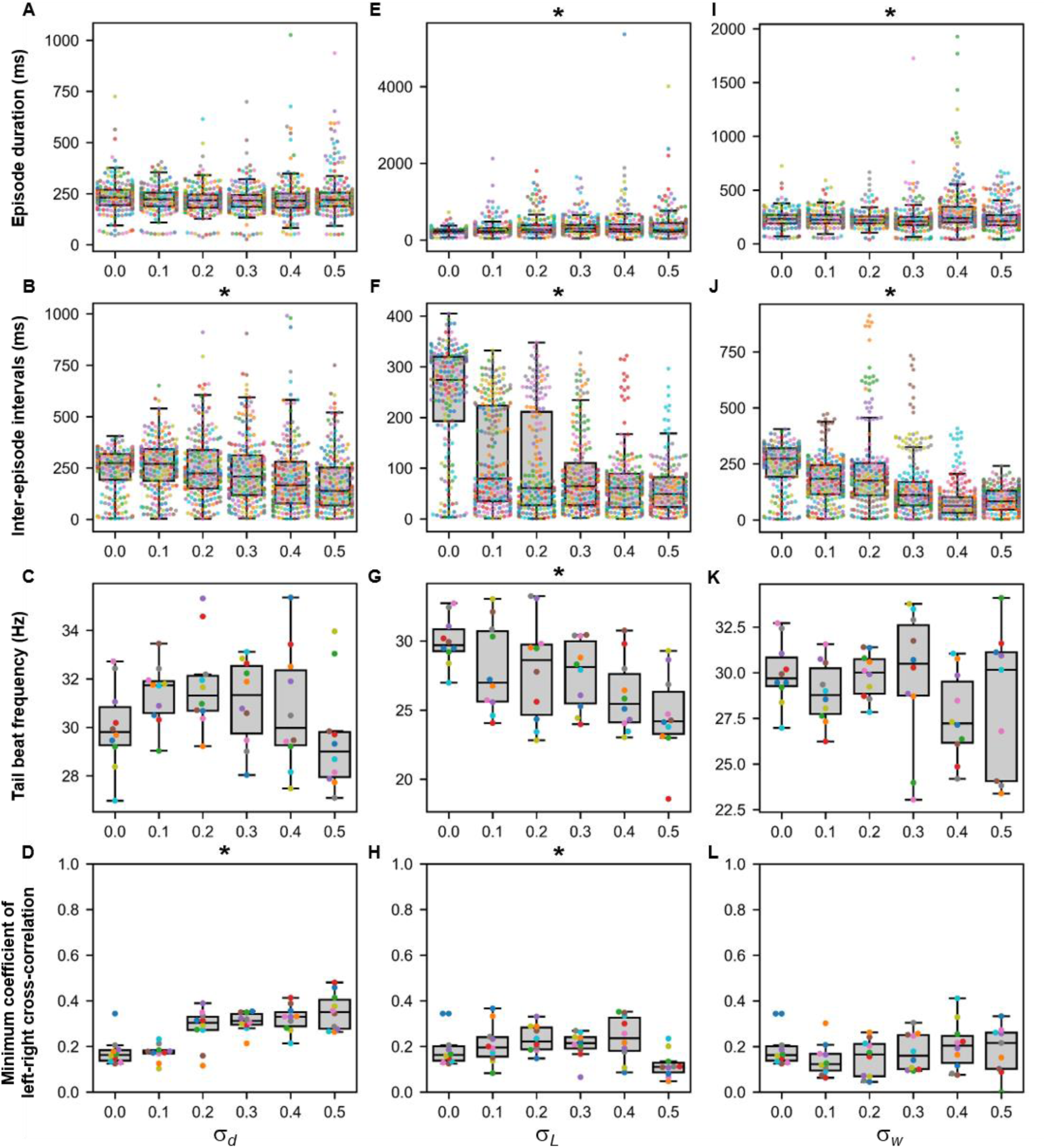
Sensitivity of beat-and-glide swimming to tonic motor command amplitude, length of rostrocaudal projections, and synaptic weighting. Ten 10,000-ms long simulations were run for each value of *σ*_*d*_ (A*-*D), *σ*_*L*_ (**E-H**), and *σ*_*w*_(**I-L**) tested*. (*A, E, I) Episode duration. (**B**, **F, J)** Inter-episode interval. (**C**, **G, K**) Average tail beat frequency during each swimming episode. (**D**, **H, L**) Minimum coefficient of the cross-correlation of left and right muscle. The minimum was taken between -10 and 10 ms time delays. Each circle represents a single swimming episode (**A**, **E, I**), inter-episode interval (**B**, **F, J**), or a single run (all other panels). Each run is color coded. Runs with only one side showing activity are not depicted in (**D**) and (**H**). Asterisks denote significant differences detected using a one-factor ANOVA test. *Statistics*: (**A**) *F*_5,1253_ = 2.5, p = 0.03. (Note that there were no pairwise differences detected). (**B**) *F*_5,1253_ = 11.2, p = 1.3 x 10^-10^. (**C**) *F*_5,54_ = 1.9, p = 0.11. (**D**) *F*_5,54_ =14.5, p = 5.2 x 10^-9^. (**E**) *F*_5,1253_ = 8.7, p = 3.8 x 10^-8^. (**F**) *F*_5,1253_ = 118.1, p = 2.0 x 10^-102^. (**G**) *F*_5,54_ = 4.0, p = 0.004. (**H**) *F*_5,54_ =3.2, p = 0.014. (**I**) *F*_5,1400_ = 13.5, p = 6.8 x 10^-13^. (**J**) *F*_5,1400_ = 74.5, p = 2.5 x 10^-69^. (**K**) *F*_5,53_ = 1.3, p = 0.30. (L) *F*_5,53_ = 0.8, p = 0.55. P- values for t-tests are found in ***Figure 9 – source data 1*.**

**Figure 10.**
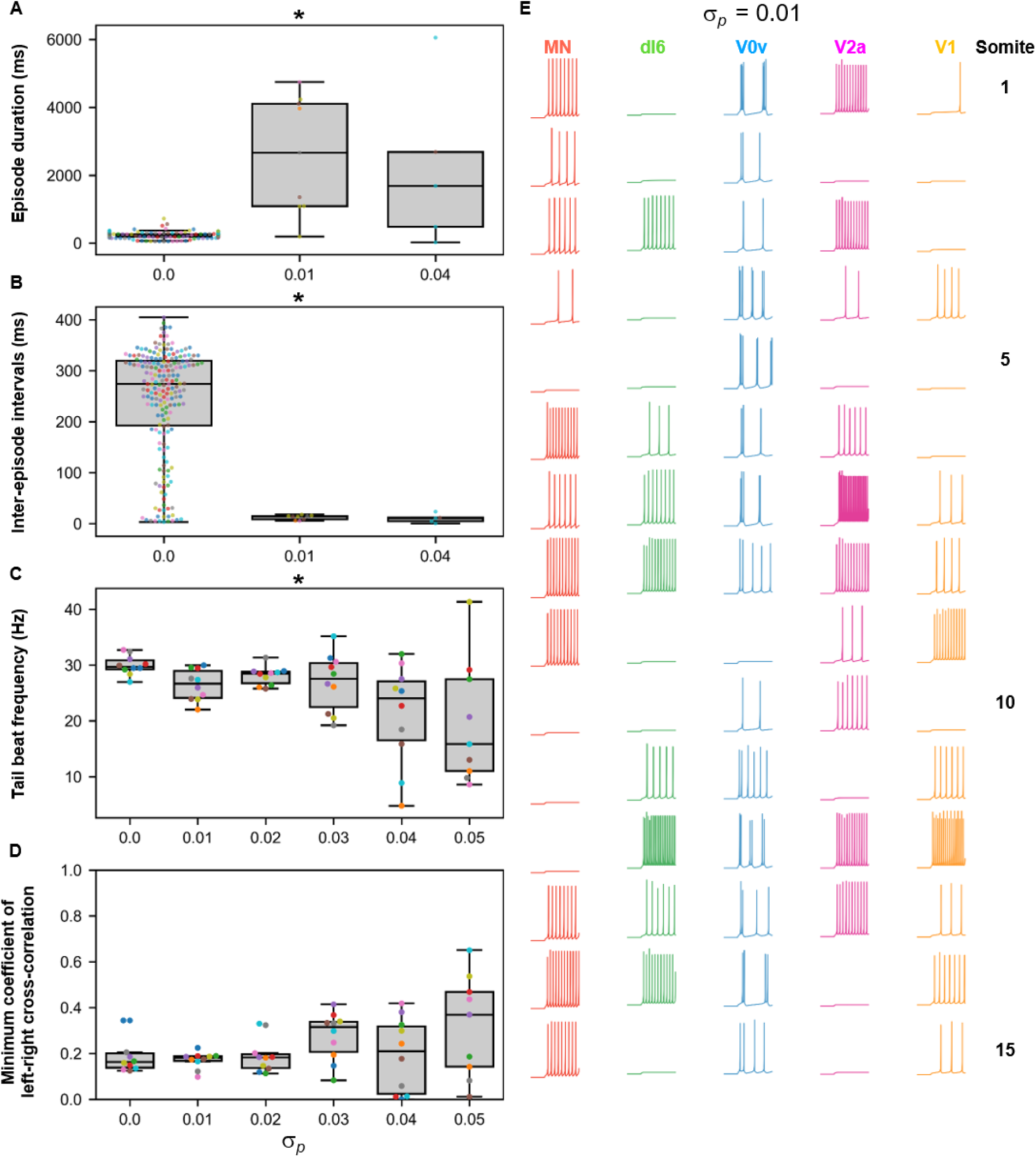
Sensitivity of beat-and-glide swimming to variability in membrane potential dynamics. Ten 10,000-ms long simulations were run at each value of *σ*_*p*_ **(A*-*D)**. (**A**) Episode duration. (**B)** Inter-episode interval. (**C**) Average tail beat frequency during each swimming episode. (**D**) Minimum coefficient of the cross-correlation of left and right muscle. The minimum was taken between -10 and 10 ms time delays. Each circle represents a single swimming episode (**A**), inter-episode interval (**B**) or a single run (**C**, **D**). Each run is color-coded. Runs not depicted exhibited either continual motor activity with no gliding pauses or no swimming activity. Asterisks denote significant differences detected using a one-factor ANOVA test. (**E**) Responses to a 1-s long step current of all neurons on the left side in a model where *σ*_*p*_= 0.01. Step current amplitudes varied between populations of neurons. The amplitude of the step currents to each population is the same as in Figure 4B. The simulation of the model with these neurons generated continued swimming activity with no gliding pauses. The neurons are ordered by somite, from somite 1 at the top to somite 15 at the bottom. ***Statistics***: (**A**) *F*_2,211_ = 143.8, p = 4.0 x 10^-40^. (**B**) *F*_2,211_ = 32.3, p = 5.8 x 10^-13^. (**C**) *F*_5,53_ = 4.0, p = 0.0036. (**D**) *F*_5,53_ = 2.1, p = 0.085. P-values for t-tests are found in ***Figure 10* – *source data 1*.**

Smaller variability in the parameters shaping the membrane potential dynamics (*σ*_*p*_) disrupted the beat-and-glide pattern with no beat-and-glide swimming observed at some values of *σ*_*p*_ (**Figure 10*A*,*B***). As *σ*_*p*_ was increased, the episode duration, inter-episode interval, and tail beat frequency were disrupted (**Figure 10*C,D***). Even slight variability in the parameters shaping membrane potential dynamics (e.g. *σ*_*p*_ = 0.01) resulted in changes in membrane excitability and, in some cases, firing patterns as evidenced by the conversion of some V0vs from burst to tonic firing (**Figure 10*E***). Thus, the beat-and-glide model was most susceptible to variations in the parameters determining membrane potential dynamics and similarly sensitive to the other parameters tested.

## DISCUSSION

To our knowledge, this study presents some of the first models of spinal locomotor circuits in developing zebrafish. We have built several spinal locomotor circuit models that generate locomotor movements of the developing zebrafish (**Figure 11**). These models support mechanisms of network operation of developing zebrafish spinal locomotor circuits described experimentally. Our models suggest that the circuitry driving locomotor movements could switch from a pacemaker kernel located rostrally during coiling maneuvers to network-based spinal circuits during swimming. Results from simulations where populations of spinal neurons are silenced were consistent with experimental studies. Our sensitivity analysis suggests that the correct operation of spinal circuits for locomotion is not immune to variations in firing behaviours, length of axonal projections, motor command amplitude, and synaptic weighting. The sensitivity to these parameters is variable, however.

**Figure 11.**
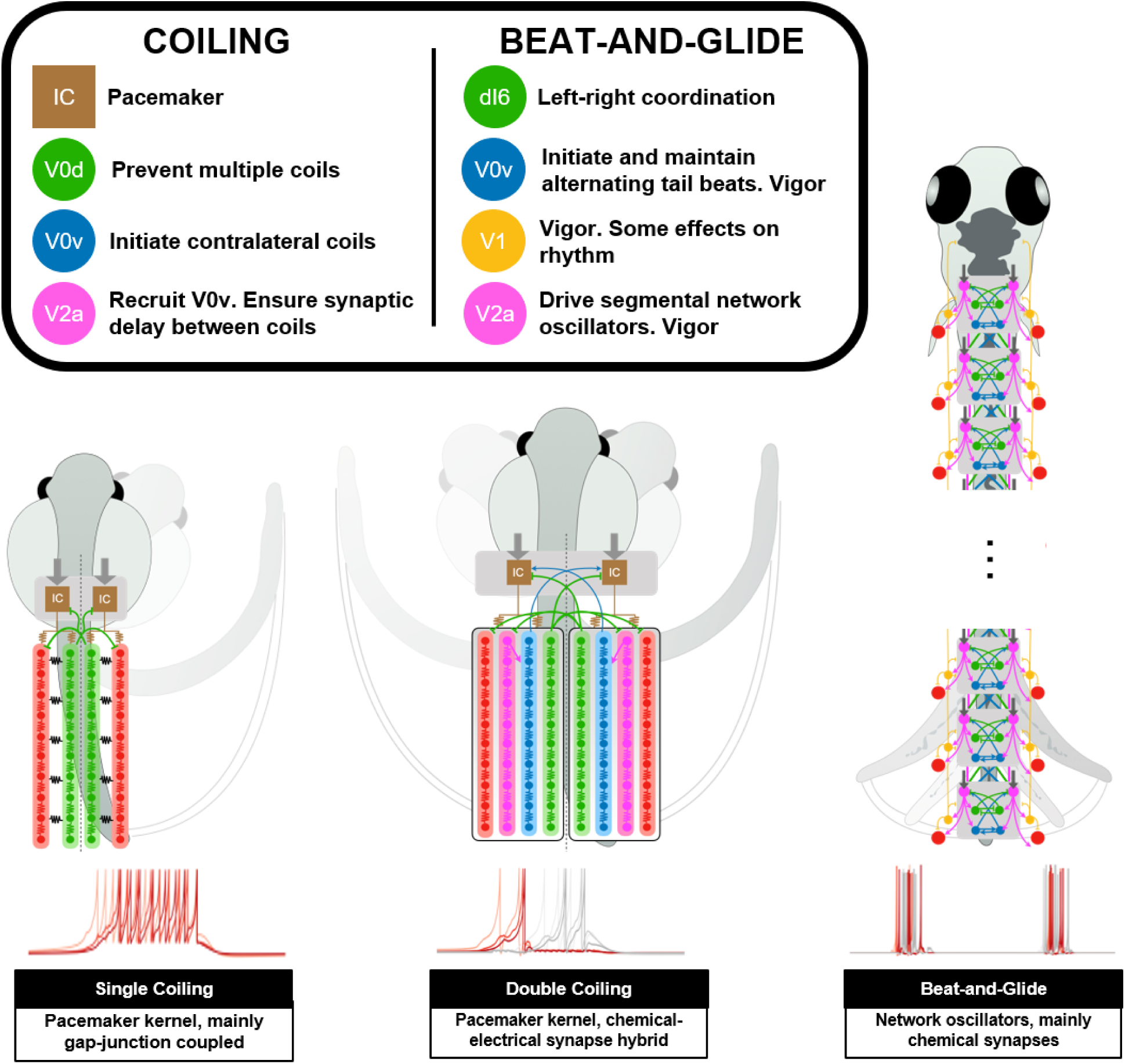
Summary figure of computational models of zebrafish locomotor movements during development.

### Pacemaker-based network for early behaviours

The earliest locomotor behaviours in zebrafish, namely single and multiple coilings, require global recruitments of neurons to synchronously contract all ipsilateral muscles (Warp et al., 2012). Electrical coupling, which lacks the delays inherent with chemical neurotransmission, enables these types of ballistic movements. Early locomotor behaviour in zebrafish seems to rely on this architecture, as demonstrated by the necessity of electrical but not chemical synapses (Saint-Amant & Drapeau, 2000, 2001). The rapid and multidirectional current transmission supported by electrical synapses is a perfect solution for en masse activation of a neural circuit (Bennett & Zukin, 2004). However, synchronous activation of an ensemble of neurons does not accommodate rhythmic activity, which requires more precise timing and connection strength. For example, the emergence of double coiling in our model was generated by chemical synaptic excitation of contralateral pacemaker neurons that had to be sufficiently delayed to enable the first coil to complete before initiating the second contralateral coil. Commissural glycinergic transmission was also required to tamper down coiling events with more than two successive coils. Suppose multiple coiling is a preparatory stage towards the emergence of repetitive, left-right alternating tail beats. In that case, the possible importance of contralateral excitation and inhibition at this stage presages the establishment of similar operational mechanisms to the generation of swimming.

### Network oscillators for swimming movements

To generate swimming, we delegated the generation of the rhythm driving tail beats to network oscillators distributed along the length of the spinal cord. Spinal locomotor circuits may transition away from pacemakers as the source of the rhythm to prevent being vulnerable to any flaws in the function of a small population of neurons. Also, there may be multiple local rhythms that control body oscillations along the developing zebrafish’s length. Indeed, locomotor output has proven to be very robust to the sectioning of the spinal cord, leading to the suggestion that redundant rhythm-generating circuits must be present within the spinal cord (McDearmid & Drapeau, 2006; Wiggin et al., 2012, 2014). Experimental evidence from our lab further suggests that a transition from a rhythm driven by a pacemaker kernel to a rhythm driven by local network oscillators occurs progressively from the caudal toward the rostral end of the body (Roussel et al., 2020).

The V2as are well recognized as the neural engine that drives swimming activity in zebrafish spinal circuits (Eklof-Ljunggren et al., 2012; Ljunggren et al., 2014). While some V2a interneurons have shown intrinsic burst firing in the adult zebrafish (Song et al., 2018, 2020), V2a interneurons in developing zebrafish show either tonic or modestly spike adapting firing (Menelaou & McLean, 2019). We thus sought to generate beat-and-glide swimming with tonically firing V2as. Successive left-right alternating tail beats were generated by combining contralateral excitation from bursting commissural excitatory neurons to initiate alternating tail beats and contralateral inhibition to prevent co-contraction of both sides. In fact, a simulation with only tonic firing neurons could also generate beat-and-glide swimming over several seconds. Thus, V2as could very well drive rhythmic tail beats in larval zebrafish while firing tonically. If this is the case, then the central role of V2as depends less on their ability to produce a bursting rhythm. Instead, the pivotal role of V2as in enabling swimming activity would be to coordinate the many spinal interneuron populations that generate the patterns of repetitive, left-right alternating tail beats seen in developing zebrafish swimming (Saint-Amant, 2010). The observation that in our beat-and-glide simulation, V2a neuron firing phasically precedes firing of all the other intrasegmental spinal interneurons and motoneurons reinforces the central role of these neurons in driving rhythmic tail beats.

We did find that in simulations where there were only tonic firing neurons, the stability of swimming episode durations started degrading after about 6,000 ms. Therefore, burst firing neurons may help to promote the stability of the beat-and-glide pattern. Whether or not this is the case remains to be tested experimentally. Neuromodulation may serve as a mechanism that permits V2as, or other spinal neurons, to toggle between tonic and burst firing through neuromodulation. It is well established that neuromodulators shape the activity of spinal locomotor circuits, likely by regulating intrinsic properties of spinal neurons and through modulation of synaptic weighting and other mechanisms. Blocking D4 dopamine receptors at 3 dpf prevents the transition from burst to beat-and-glide swimming (Lambert et al., 2012), suggesting that dopamine from supraspinal sources plays a role in setting the beat-and-glide phenotype by shortening swimming episode duration. Paired recordings of diencephalospinal dopaminergic neurons and spinal motoneurons during swimming show that these two populations often burst together (Jay et al., 2015). Later in development at 6-7 dpf, activation of D1 dopamine receptors increases the recruitment of slow motoneurons to increase swimming speed (Jha & Thirumalai, 2020). The neuromodulator serotonin (5-HT) has been found to either increase motor output by decreasing inter-episode intervals in intact larval zebrafish (Brustein, 2005; Brustein et al., 2003) or decrease swimming frequency or burst firing in spinalized larvae and adult zebrafish (Gabriel et al., 2009; Montgomery et al., 2018). In the adult zebrafish, serotonin strengthens inhibition to motoneurons between tail beats and slows down the onset of the depolarization that initiates each successive tail beat (Gabriel et al., 2009). Our model could identify possible targets within the spinal cord for specific neuromodulators of locomotor function in zebrafish.

### Modelling considerations

Our sensitivity analysis suggests that the neuromodulation of intrinsic properties that affect the membrane potential dynamics of spinal neurons could easily modulate locomotor output. The behaviour of our models was also sensitive to a lesser degree to increasing variability in descending drive, synaptic weighting, and rostrocaudal extent of connections. Variability in these parameters could change the proportions of coiling types or the values of the characteristics of swimming output measured (e.g. episode duration, inter-episode interval). Model parameter variability sometimes increased the variability of motor output (e.g. **Figure 10**), perhaps indicating a breakdown of the model. However, variability in both model parameters and motor output should not necessarily be considered weaknesses of the model but may instead reflect true biological variability (Marder & Taylor, 2011). For instance, recordings of swimming characteristics such as episode duration and inter-episode intervals in larval zebrafish show appreciable variation (Brustein, 2005; Buss & Drapeau, 2001). Quantifying heterogeneity within and between animals may guide the appropriate levels of parameter variability to include in future iterations of our models.

Many computational models have already been made of spinal circuits for swimming in species that use undulatory movements spreading from head to tail. These include models for *Xenopus* (Ferrario et al., 2018; Hull et al., 2016), lamprey (A. Kozlov et al., 2009; A. K. Kozlov et al., 2014; Messina et al., 2017), and salamanders (Bicanski et al., 2013; Ijspeert et al., 2007). These models have become detailed enough to include many neurons forming circuits distributed across the hindbrain and the spinal cord. Some models incorporate specific intrinsic and ligand-gated currents with known roles in rhythmogenesis in their respective species (Ferrario et al., 2018; A. Kozlov et al., 2009; A. K. Kozlov et al., 2014). Simulations of the models have been used to test aspects of swimming control, including steering commands from descending commands to spinal networks (A. K. Kozlov et al., 2014), the integration of sensory triggers of flocomotion (Ferrario et al., 2018; Ijspeert et al., 2007), the coupling of axial and limb central pattern generators (Ijspeert et al., 2007), and the role of left-right coupling in rhythm generation (Messina et al., 2017). Our model could be used to identify possible similarities or differences in how these aspects of motor control are controlled in the zebrafish.

### Testable Predictions

To the best of our knowledge, this is the first model to generate several forms of locomotor movements in developing zebrafish based upon previously described neurons and their connectivity patterns. The analysis of the simulations generated yielded several predictions about possible connections between spinal neurons, firing properties of neurons, and roles for neurons in specific locomotor movements. For instance, the single coiling model predicts that IC and V0d are coupled together to facilitate the activation of V0ds, which are responsible for the glycinergic synaptic bursts observed in spinal neurons at this stage (Saint-Amant & Drapeau, 2001; Tong & McDearmid, 2012).

Our modelling study also predicts that the generation of double and even multiple coils depend on untested connections between V2a to V0v neurons and between V0v to IC neurons. The latter connection would be needed to initiate consecutive left-right alternating coils through the activation of contralateral IC neurons, while the former connection would be needed to activate the ipsilateral V0v responsible for the activation of contralateral ICs. The V2a to V0v connection could be deemed unnecessary in light of possible gap junction coupling between ipsilateral IC and V0vs. However, our modelling suggests that delayed activation of V0vs would allow the ipsilateral coil to complete before activating the contralateral coil. This delay would not be possible with gap junction mediated excitation of V0vs by ipsilateral ICs. Our double-coiling model also predicts that contralateral inhibition of ICs by V0ds prevents the generation of multiple coilings. Several of these predictions are supported by pharmacological experiments suggesting that blocking glutamatergic transmission in embryonic zebrafish precludes double coiling while blocking glycinergic transmission at that stage promotes multiple coilings (Knogler et al., 2014).

The beat-and-glide model also proposes a prominent role of delayed contralateral excitation in ensuring repetitive left-right alternating tail beats during swimming. Whether delayed contralateral excitation is a conserved mechanism of operation in double coiling and swimming remains to be tested experimentally. While V0v neurons are the likely candidate to mediate the activation of contralateral movements, different subgroups of V0v neurons are probably involved in coiling versus swimming (Björnfors & El Manira, 2016; Jay & McLean, 2019) considering the two different targets of contralateral excitation involved, namely ICs in coiling and V2as in swimming. The continued presence of left-right tail beats in simulations where the dI6 population of commissural inhibitory neurons were silenced or in simulations with blockade of glycinergic transmission further underscores the need to test the contributions of V0v neurons to left-right alternation.

Finally, the ability of our model to generate beat-and-glide swimming with or without burst firing neurons suggests a possible degeneracy in the operation of spinal swimming circuits of the developing zebrafish. This possibility would be consistent with the well-characterized degeneracy of the nervous system, as reinforced by modelling studies where combinations of intrinsic properties or connectivity can generate the same motor output (Goldman et al., 2001; Taylor et al., 2009). Many rhythmogenic currents (e.g. NMDA, calcium-dependent potassium currents, persistent sodium currents) have been implicated in the operation of locomotor circuits of zebrafish (Song et al., 2020) and other invertebrate and vertebrate rhythm-generating circuits (Anderson et al., 2012; Golowasch & Marder, 1992; Ryczko et al., 2010; Tazerart et al., 2007; Zhong et al., 2007). In addition, while some motor systems rely upon pacemaker neurons, other rhythmic motor systems could also rely on network-based mechanisms (Del Negro et al., 2010), further demonstrating the diversity of means by which the nervous system generates rhythmic activity. Whether the spinal circuits for swimming are degenerate or degeneracy is only exhibited in our modelling remains to be tested experimentally. The operation of the spinal swimming circuit in zebrafish may exhibit degeneracy dependent upon specific environmental or physiological conditions (Vogelstein et al., 2014) and their resulting neuromodulatory states.

### Future directions

Our models will require integrating additional cell populations and circuitry to capture the full range of locomotor movements of developing zebrafish. The beat-and-glide model only generates swimming within a narrow frequency range. The generation of a broader range of swimming frequency (McLean & Fetcho, 2009) will require expanding each cell population into subgroups with different intrinsic properties (Menelaou & McLean, 2012; Song et al., 2018), rostrocaudal projection patterns, and specific connectivity patterns between subgroups and between cell populations (Ampatzis et al., 2014; Bagnall & McLean, 2014; Kimura & Higashijima, 2019; Menelaou & McLean, 2019; Sengupta et al., 2021; Song et al., 2020). These subgroups, which may arise from different birth dates (McLean & Fetcho, 2009; Satou et al., 2012), are active at specific swimming frequencies (McLean et al., 2007, 2008; McLean & Fetcho, 2009). There seem to be modules consisting of neurons within each cell population that are active at specific swim frequencies (Ampatzis et al., 2014; Menelaou & McLean, 2019; Song et al., 2018, 2020). Indeed, previous studies in zebrafish have shown that MNs and V2a neurons are organized in three different modules (linked to slow, medium, and fast MNs) that are differentially recruited according to swim frequency (Ampatzis et al., 2014; Song et al., 2020). Swim frequency modules likely include commissural excitatory V0v interneurons (Björnfors & El Manira, 2016; McLean et al., 2008) and commissural inhibitory interneurons belonging to either the dI6 or V0d populations (Satou et al., 2020). The modelling of additional subgroups, especially in the context of swim-frequency modules, will need to take into account the high specificity of connectivity between subgroups within a cell population (Menelaou & McLean, 2019; Song et al., 2020) and subgroups belonging to different spinal populations within swim frequency-modules (Ampatzis et al., 2014; Bagnall & McLean, 2014; Menelaou & McLean, 2019; Song et al., 2020).

Subgroups within cell populations are not necessarily restricted to those belonging to different swim frequency modules but may also exist between neurons involved in rhythm versus vigor of movement. Subgroups for vigor seem to be present within the V2a (Menelaou & McLean, 2019; Song et al., 2018) and V0v (Björnfors & El Manira, 2016; Jay & McLean, 2019; McLean et al., 2007) populations. Furthermore, the implementation of circuitry for swimming vigor is likely to necessitate adding the ipsilaterally projecting, inhibitory V2b population (Callahan et al., 2019). The circuits for frequency and vigor are likely to interact, as seen by the swimming frequency-dependent action of V1 neurons (Kimura & Higashijima, 2019). Frequency and vigor are also likely to be shaped by sensory information. Incorporating spinal neurons that integrate sensory information (Y. C. Liu & Hale, 2017) provided by peripherally-located and spinally-located sensory neurons (Böhm et al., 2016; Picton et al., 2021) will provide a more accurate representation of swimming control at the level of the spinal cord.

Finally, the undefined role of specific spinal neuron populations could be studied after being integrated into the model following further characterization. For example, ventral V3 neurons in mouse spinal locomotor networks have been studied using modelling. Those studies suggest an important role for these neurons in left-right coordination in mouse locomotion (Danner et al., 2019). Similar computational studies using our model could reveal testable predictions of the role of these neurons (England et al., 2011; Yang et al., 2010) in zebrafish swimming.

Our models simulate several developmental milestones of the zebrafish locomotor behaviour. Iterative changes were made to each model to successively transition from single coiling to double coiling and then to beat-and-glide swimming. This iterative process could be further developed to obtain a higher resolution understanding of the maturation of locomotion in zebrafish. Further transitory models could be built to fill the gaps between our current models (e.g. a model for burst swimming that precedes beat-and-glide swimming). The generation of these additional transitory models could be coupled with experimental data studying the mechanisms that drive the transition from one milestone to the other (Brustein, 2005; Knogler et al., 2014; Lambert et al., 2012; Roussel et al., 2020) to identify specific underlying changes in intrinsic and network properties. Thus, the models presented herein offer invaluable tools to investigate further the mechanisms by which spinal circuits control facets of swimming, including speed, direction, and intensity through interactions within the spinal cord and with supraspinal command centres, as well as the developmental dynamics that ensure proper maturation of movement during development.

## MATERIALS AND METHODS

### Modelling environment

Modelling was performed using Python 3.6.3 64-bits. We did not analyze the early parts of simulations (up to 200 ms) to allow the effects of initial conditions to dissipate.

### Modelling of single neurons

We modelled neurons using a single compartment, simple spiking neuron model developed by Izhikevich (2007). The following general differential equations govern the dynamics of the membrane potential:

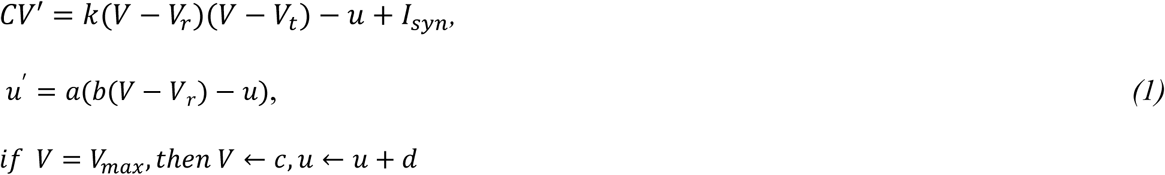

Specific active conductances are not included in these models. Instead, values of the parameters *a, b, c*, *d,* and *V_max_* (which respectively represent the time scale of the recovery variable *u*, the sensitivity of *u* to the sub-threshold variation of *V*, the reset value of *V* after a spike, the reset value of *u*, and the action potential peak), as well as values of the parameters *k*, *C*, *V_r_*, and *V_t_* (coefficient for the approximation of the subthreshold part of the fast component of the current-voltage relationship of the neuron, cell capacitance, resting membrane potential, and threshold of action potential firing) can be selected to model a wide range of firing behaviours including bursting (or chattering) pacemaker, tonic firing, phasic spiking neurons or firing rate adaptation neurons **(Table 2)**. *I_syn_* represents the sum of the synaptic and gap junction currents received by the neuron. For all models, the Euler method was used for solving ordinary differential equations with a time step of 0.1 ms.

**Table 2.**
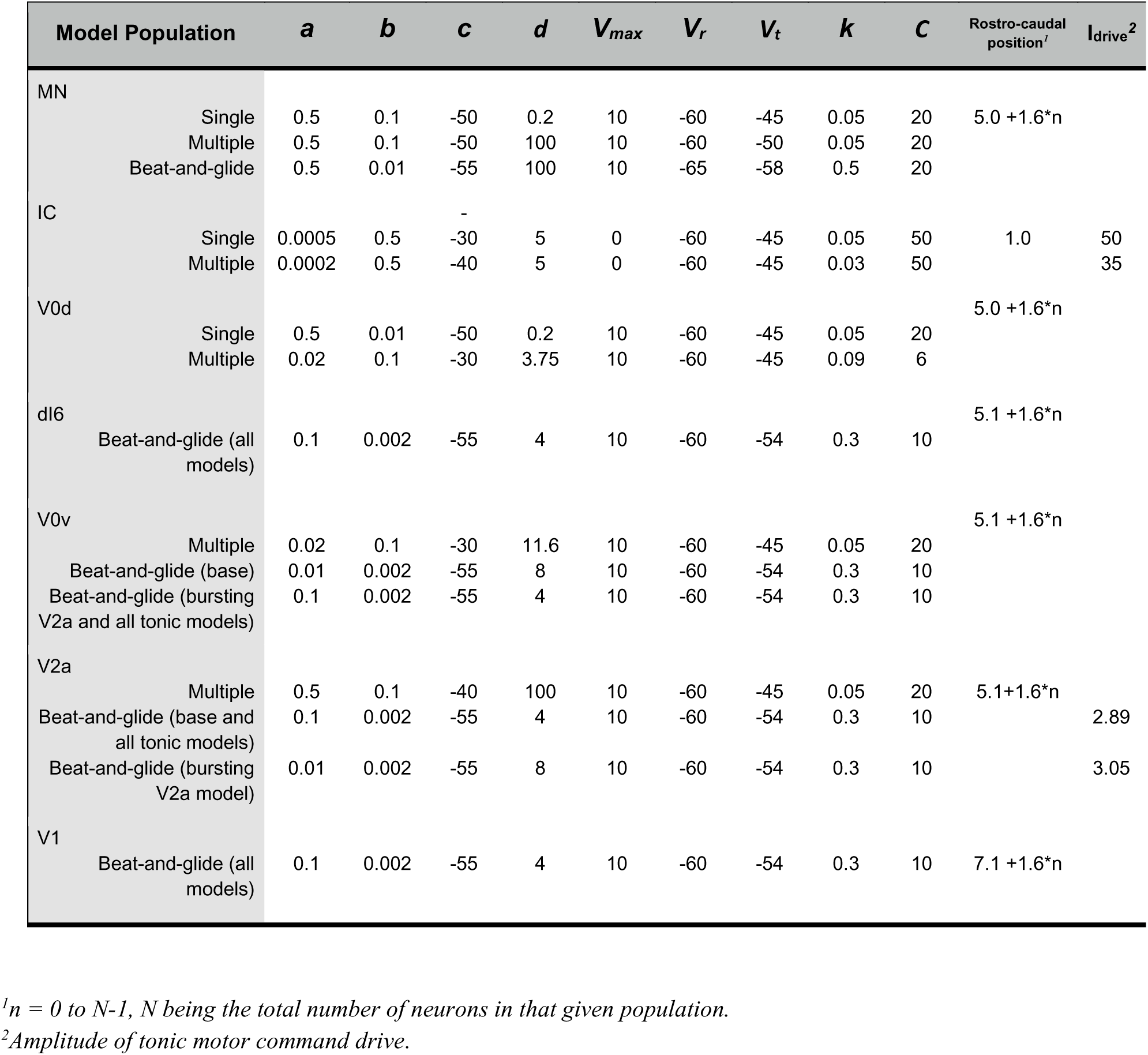
Parameter values of neurons

### Modelling synapses

We modelled all electrical synapses (i.e. gap junctions) as ideal resistors following Ohm’s Law:

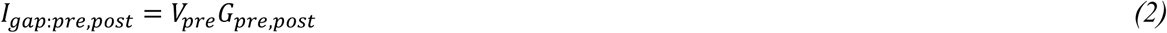

With *I_gap:pre,post_* representing the synaptic current flowing to the postsynaptic neuron from the presynaptic neuron through gap junctions and *G_pre,post_* the total conductance of gap junctions between the presynaptic and postsynaptic neurons **(Table 3)**.

**Table 3.** Electrical synapse (gap junctions) weights between neuron populations.

| Coiling<br>Beat-and-glide | MN | IC | V0d<br>dl6 | V0v | V2a |
| --- | --- | --- | --- | --- | --- |
| **MN** |  |  |  |  |  |
| Single-coiling | 0.1 |  |  |  |  |
| Double-coiling | 0.07 |  |  |  |  |
| Beat-and-glide (all<br>models) | 0.005 |  |  |  |  |
| **IC** |  |  |  |  |  |
| Single-coiling | 0.04 | 0.001 |  |  |  |
| Double-coiling | 0.03 | 0.0001 |  |  |  |
| **V0d or dl6** |  |  |  |  |  |
| Single-coiling | 0.01 | 0.05 | 0.04 |  |  |
| Double-coiling | 0.0001 | 0.05 | 0.04 |  |  |
| Beat-and-glide (all<br>models) | 0.0001 |  | 0.04 |  |  |
| **V0v** |  |  |  |  |  |
| Double-coiling | 0.0001 | 0.0005 |  | 0.05 |  |
| Beat-and-glide (all<br>models) | 0.005 |  |  | 0.05 |  |
| **V2a** |  |  |  |  |  |
| Double-coiling | 0.005 | 0.15 |  |  | 0.005 |
| Beat-and-glide (all<br>models) | 0.005 |  |  |  | 0.005 |

Synaptic conductances of chemical synapses were modelled as a sum of two exponentials weighted by a synaptic weight based upon the general equation:

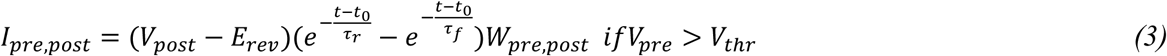

Where *I_pre,post_* is the synaptic current received by the postsynaptic neuron from neurotransmitter release by the presynaptic neuron if the presynaptic neuron membrane potential, *V_pre_*, crosses a voltage threshold, *V_thr_*, at the synapse. *V_post_* is the membrane potential of the postsynaptic neuron, *E_rev_* is the reversal potential, *τ_r_* and *τ_f_* are the rise and fall time constants, respectively, *t_0_* is the time at which *V_pre_* crossed *V_thr_*, and *W_pre,post_* is the synaptic weight between the presynaptic and postsynaptic neurons (**Table 4**). *I_pre,post_* is equal to 0 if *V_pre_* is below *V_thr_*.

**Table 4.** Chemical synapse weights between neuron populations. Pre-synaptic neurons are in rows. Post-synaptic neurons in columns.

| Pre-synaptic | Post-synaptic |  |  |  |  |  |  |
| --- | --- | --- | --- | --- | --- | --- | --- |
|  | MN | IC | V0d<br>dl6 | V0v | V2a | V1 | Muscle |
| **MN** |  |  |  |  |  |  |  |
| Single coiling |  |  |  |  |  |  | 0.015 |
| Double coiling |  |  |  |  |  |  | 0.02 |
| Beat-and-glide (all models) |  |  |  |  |  |  | 0.1 |
| **V0d** |  |  |  |  |  |  |  |
| Single coiling | 0.3 | 0.3 |  |  |  |  |  |
| Double coiling | 2.0 | 2.0 |  |  | 2.0 |  |  |
| **dl6** |  |  |  |  |  |  |  |
| Beat-and-glide (base) | 1.5 |  | 0.25* |  | 1.5 |  |  |
| Beat-and-glide (bursting V2a) | 1.5 |  | 0.25* |  | 2.0 |  |  |
| Beat-and-glide (all tonic neurons) | 1.5 |  | 0.25* |  | 1.5 |  |  |
| **V0v** |  |  |  |  |  |  |  |
| Double coiling |  | 0.175 |  |  |  |  |  |
| Beat-and-glide (base) |  |  |  |  | 0.4 |  |  |
| Beat-and-glide (bursting V2a) |  |  |  |  | 0.75 |  |  |
| Beat-and-glide (all tonic neurons) |  |  |  |  | 0.4 |  |  |
| **V2a** |  |  |  |  |  |  |  |
| Double coiling |  |  |  | 0.04 |  |  |  |
| Beat-and-glide (base) | 0.5 |  | 0.5 | 0.3 | 0.3 | 0.5 |  |
| Beat-and-glide (bursting V2a) | 0.5 |  | 0.75 | 0.275 | 0.3 | 0.5 |  |
| Beat-and-glide (all tonic neurons) | 0.5 |  | 0.5 | 0.25 | 0.3 |  |  |
| **V1** |  |  |  |  |  |  |  |
| Beat-and-glide (base) | 1.0 |  | 0.2 | 0.1 | 0.5 |  |  |
| Beat-and-glide (bursting V2a) | 1.0 |  | 0.2 | 0.1 | 0.5 |  |  |
| Beat-and-glide (all tonic neurons) | 1.0 |  | 0.2 | 0.1 | 0.6 |  |  |

\*Scaled by a random number selected from a gaussian distribution with mean of 1 and variance of 0.1.

We implemented two types of chemical synapses: glutamatergic and glycinergic synapses. The former differs from the latter by the respective reversal potential values *E_rev_* of glutamatergic and glycinergic synapses and the time constant values *τ_r_* and *τ_f_* **(Table 5)**. Values of the glycinergic *E_rev_* are depolarized at early developmental stages (Saint-Amant & Drapeau, 2000, 2001), and this reversal potential becomes gradually hyperpolarized (Ben-Ari, 2002). All chemical synapses were turned off in the initial 50 ms of every simulation to allow initial conditions to dissipate.

**Table 5.** Glutamatergic and glycinergic reversal potentials and time constants

| Chemical Synapse | $E_{rev}$ | $\tau_r$ | $\tau_f$ | $V_{thr}$ |
| --- | --- | --- | --- | --- |
| Glutamatergic | 0 | 0.5 | 1.0 | -15 |
| Glycinergic | -45, -58, -70* | 0.5 | 1.0 | -15 |

\* *for single and double coiling and beat-and-glide swimming models, respectively.*

### Spatial arrangement of spinal neurons

A key feature of our modelling approach was to assign spatial coordinates *(x, y)* to point-like neurons (i.e. neurons have no spatial dimension, but they have a position in space), giving the spatial distribution of neurons a central place in our model computing process. We used the Euclidean distance to calculate the distance between each neuron and to approximate axon length. Distance unit is arbitrary and was set so that one model somite was 1.6 arbitrary distance units (a.d.u.) long. Time delays for each synaptic connection were computed as a function of the distance between neurons and were used to calculate delayed synaptic current:

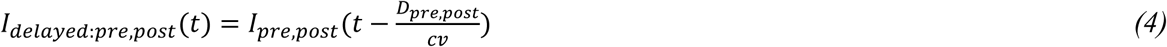

With *D_pre,post_* as the Euclidean distance between the presynaptic and postsynaptic neurons and *cv* as the transmission speed in arbitrary distance units per second (a.d.u./s). This distance and the neuron position were also used to apply conditions on synaptic weights of neurons (e.g. limits as to how far descending neurons project). For the single coiling model, *cv* was set to 4.0 a.d.u/s. For the multiple coiling model, *cv* was set to 1.0 a.d.u/s. These values were obtained through trial-and-error and may reflect changes in myelination and body size of the developing zebrafish. For the beat-and-glide swimming model, *cv* was set to 0.8 a.d.u/s, which led to intersegmental transmission delays in the range of 3.0-4.0 ms, closely matching the 1.6 ms intersomitic delay previously reported (McDearmid & Drapeau, 2006), assuming that each model somite represents two biological somites at this developmental stage (see *Musculoskeletal model* below).

Spinal locomotor circuits were distributed across two columns, one for each side of the body, giving the network a nearly one-dimensional organization along the rostrocaudal axis. Therefore, we used the *x*-axis as the rostrocaudal axis, whereas the *y*-axis was only used to partition neurons from the left and right sides (assigning the coordinate *y* = 1 a.d.u. for the right side and *y* = −1 a.d.u. for the left side).

### Sensitivity testing

We scaled key parameters to Gaussian noise to test the robustness of our three base models (single coiling, multiple coiling, and beat-and-glide swimming) to parameter variability. Sensitivity to noise of the base models was tested by scaling the parameters that set the tonic motor command drive’s amplitude, the rostrocaudal length of neuron projections, the membrane potential dynamics (Izhikevich model), and synaptic weighting. These four sets of parameters were randomized by multiplying the parameters with a random number picked from a Gaussian distribution with mean, µ = 1, and standard deviations, *σ*_*d*_, *σ*_*l*_, *σ*_*p*_, and *σ*_*w*_, respectively. The amplitude of the motor command drive was randomized at each time point. The parameters for the membrane potential dynamics, rostrocaudal length of axons, and synaptic weights were randomized at the start of each simulation and did not change during the simulations.

### Musculoskeletal model

We implemented a musculoskeletal model of the fish body to convert the output of the spinal circuit model into changes in body angles and frequency of locomotor movements. Each MN output along the fish body was inputted into a muscle cell (**Figure 1**). The membrane potential of the muscle (*V*) was modelled as a simple passive RC circuit (*R* and *C* being the muscle cell membrane resistance and capacitance, respectively), described by the following equation:

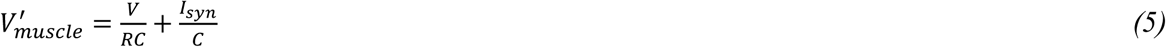

For muscle cells, values of R were 25 (single coiling), 50 (double coiling), and 1 (beat-and-glide); values of C were 10 (single coiling), 5 (double coiling), and 3 (beat-and-glide). These values were chosen to produce kinematics representative of those seen experimentally. To reduce computational load, we modelled one muscle cell as representing three somites of the body in the base model for coiling and two somites of the body in the base model for swimming. The whole body of the fish was modelled as a chain of uncoupled damped pendulums. We computed local body angles according to the difference in activity between the local left and right muscle cells. The deflection angle *θ*_*i*_ of the i^th^ muscle cell was computed according to the following differential equation.

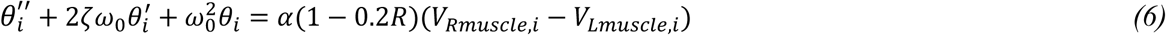

With *V*_Rmuscle,i_ and *V*_*Lmuscle,i*_ being the solution of the equation *(5)* for the *i*^th^ muscle on the right and left side of the body, respectively (**Figure 1*D***). *α* is the conversion coefficient from an electric drive of the muscle cells to a mechanical contraction of the same cells. The midline of the body can be computed at any given time as (*x*,*y*) coordinates using trigonometric identities from *θ*_*i*_ (**Figure 1*E***). Specifically,

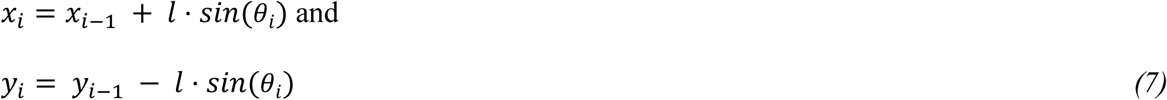

where (*x*_*i*_, *y*_*i*_) are the spatial coordinates of the *i*^th^ somite, and *l* is its length. We set (*x*_0_, *y*_0_) to (0, 0) and applied the previous set of equations *(7)* for *i* ≥ 1. Thus, heat-maps of local body angle (*θ*_*i*_) variation through time provide comprehensive information about the network output (**Figure 1*F***). The integrated motor output of the model (for example, **Figure 5*B***) was calculated as the sum of the muscle output at all muscle cells on both sides of the body, followed by a convolution of this sum with a 50 ms square wave. Left-right alternation at the *i*^th^ somite was analyzed using cross-correlation of *V*_Rmuscle,i_ and *V*_*Lmuscle,i*_ at that somite. The minimum coefficient in the range of time delays between -20 and 20 ms was calculated to estimate left-right alternation. A value of 0 indicates left-right out-of-phase alternation, while a value of 1 suggests complete in-phase synchrony.

### Analysis of locomotor activity

To calculate the duration of swimming episodes, we summated the muscle activity across all somites from both sides of the body. This muscle activity was then convoluted, and a threshold of 0.5 arbitrary units was set to detect the start and end of each swimming episode of most simulations. In a few simulations where motor output was very large, the threshold was adjusted to detect episodes. To estimate the tail beat frequency, we determined when the most caudal somite crossed the midline of the body of the musculoskeletal model (a threshold of 0.5 arbitrary units from the center was used to detect crossing to a side of the body). The reciprocal of the interval between consecutive left-to-right or right-to-left crossing was used to calculate the instantaneous tail beat frequency. Any interval greater than 100 ms was considered to be between episodes rather than within an episode and discarded from the calculation of instantaneous tail beat frequency.

To calculate the phase delay between pairs of neurons in the beat-and-glide swimming model, we first calculated the autocorrelation of the reference neuron. The time delay at which the peak autocorrelation occurred was used to estimate the period of the reference neuron cycle. The cross-correlation between the reference and test neuron was then calculated, and the phase delay was calculated as the time delay at which the peak of the cross-correlation occurred divided by the cycle period of the reference neuron in radians. In the coiling models, the cycle period is 1,000-2,000 ms (single coiling) or 10,000-20,000 ms (double coiling) due to longer inter-coiling intervals. Normalizing phase shifts by this cycle period makes the phase delays very small. Therefore, for the coiling models, the period of the reference neuron cycle was estimated by the average duration of single coiling or double coiling events. Note that this procedure does not change the polarity of the phase delay but better separates the various phase delays on a polar plot.

### Statistical Analysis

Statistical analysis was performed using the SciPy Python library. Statistical tests consisted of one-factor ANOVA tests followed by two-tailed Student’s t-tests. A p-value < 0.05 was used to determine statistical significance, and all tests were corrected for multiple comparisons (Bonferroni correction for multiple t-tests).

### Availability of code

The code for the models can be accessed at https://github.com/bui-lab/code (pending acceptance of the manuscript). Updates and revisions to the models will also be made available at this site.

## Acknowledgments

We would like to thank Martha Bagnall, Vamsi Daliparthi, Joe Fetcho, Sara Goltash, Michael Hildebrand, Alex Laliberte, Nicolas Lalonde, John Lewis, Aaron Shifman, and Emily Standen for advice related to the modelling and critical discussion of this manuscript. This research was supported by an NSERC Discovery Grant (RGPIN-2015–06403), an NSERC Canadian Graduate Scholarship M award (NSERC 712210101627) and a McDonnell Center for Cellular and Molecular Neurobiology Postdoc Fellowship FY21.

## FIGURE TITLES AND LEGENDS

**Figure 2 - figure supplement 1.**
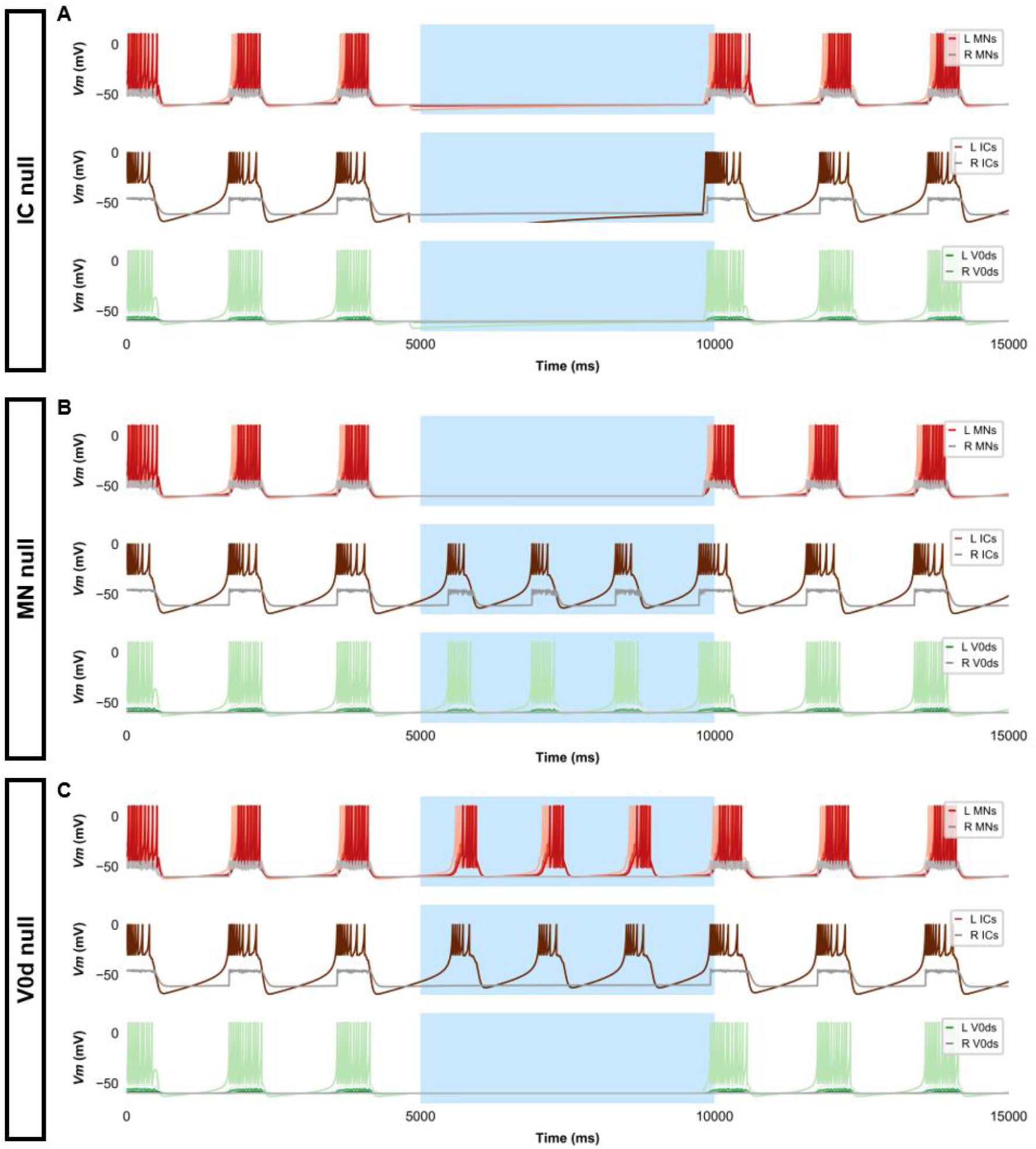
Silencing spinal neurons during single coiling. Simulations consisted of three 5,000 ms epochs. In the middle epoch, silencing of targeted spinal neurons was achieved by removing all synaptic and external currents from the targeted population. Synaptic and external currents were restored in the last epoch. (**A**) Silencing IC neurons silences the other spinal neurons. (**B**) Silencing MNs slightly reduces IC burst duration but does not preclude IC bursting. (**C**) Silencing V0ds blocks synaptic bursts in contralateral ICs and MNs but does not preclude single coils, nor does it lead to multiple coils. The *Vm* of a rostral (lightest), middle, and caudal (darkest) neuron is shown, except for IC neurons that are all in a rostral kernel. L: left, R: right.

**Figure 2 - figure supplement 2.**
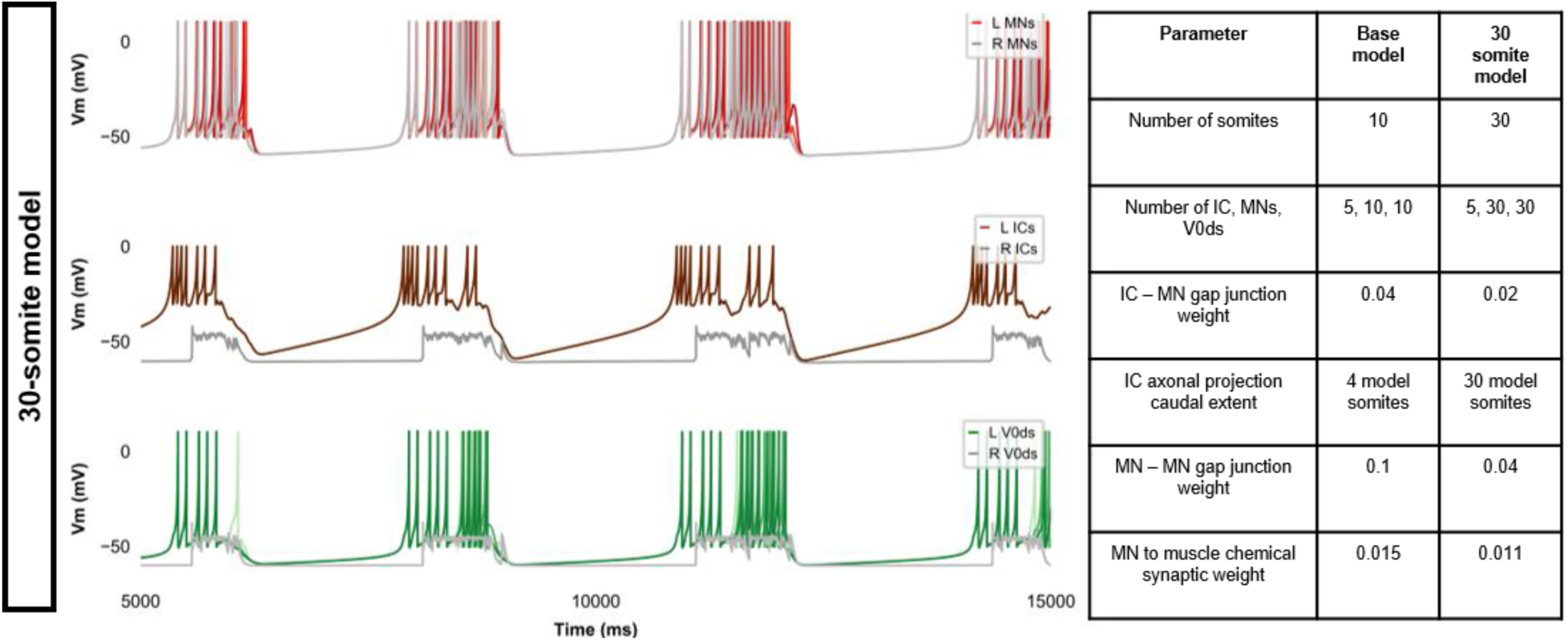
Membrane potential (*Vm*) during a simulation of a 30- somite single-coiling model. The *Vm* of a rostral (lightest), middle, and caudal (darkest) neuron is shown, except for IC neurons that are all in a rostral kernel. L: left, R: right.

**Figure 3 – figure supplement 1.**
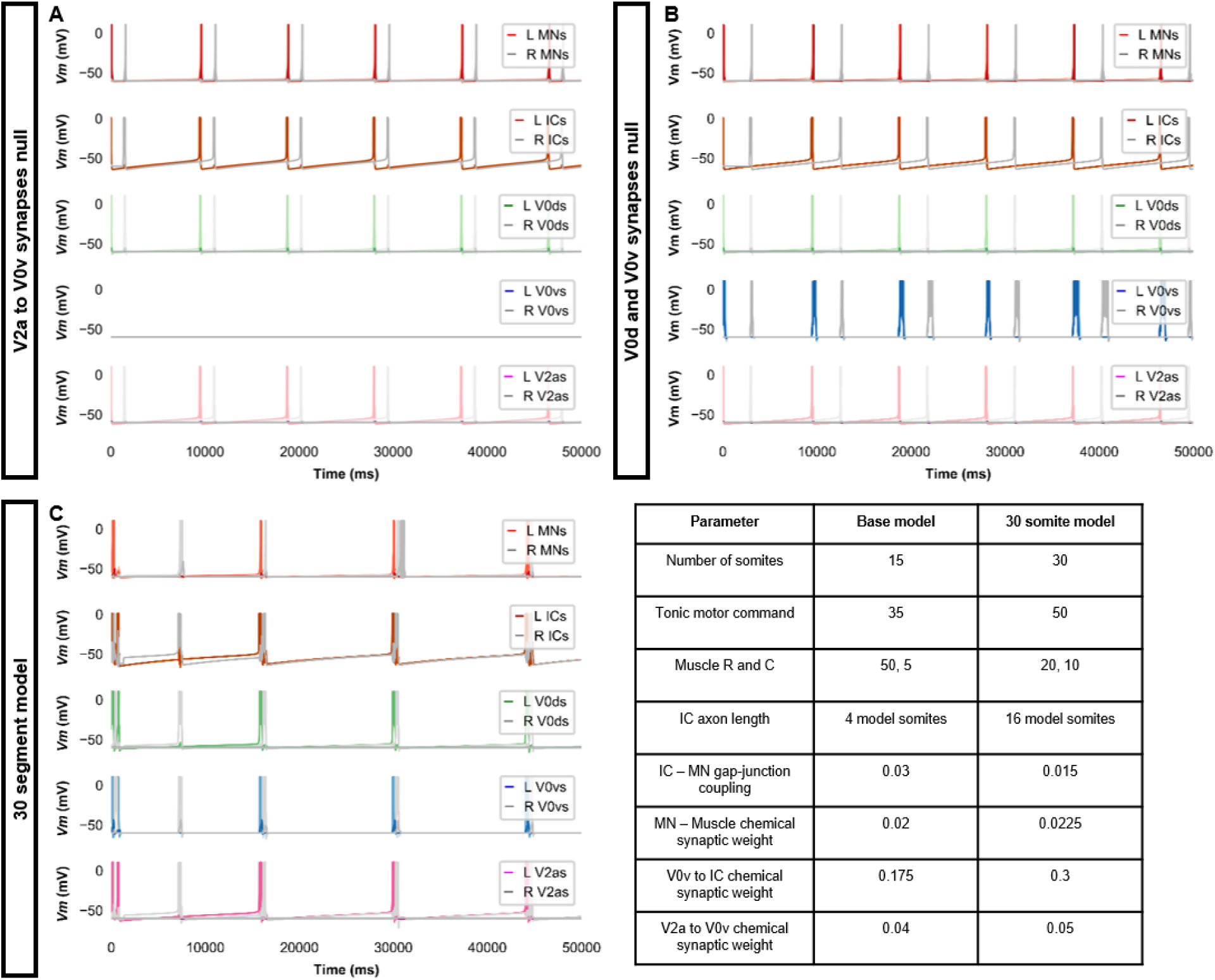
Double coiling model with no V2a to V0v synapses, no contralateral synapses, or with 30 somites. **(A)** Membrane potential (*Vm*) during a simulation without V2a to V0v synapses. V0v neurons remain inactive, and there are only single coils. **(B)** Simulation with no contralateral inhibition or excitation. The lack of double and multiple coils, even without contralateral inhibition, suggests that contralateral excitation is necessary to generate double and multiple coils. (**C**) Double coiling in a model composed of 30 somites. The *Vm* of a rostral (lightest), middle, and caudal (darkest) neuron is shown, except for IC neurons that are all in a rostral kernel. L: left, R: right. See also ***Figure 3* - *video 5***.

**Figure 3 – figure supplement 2.**
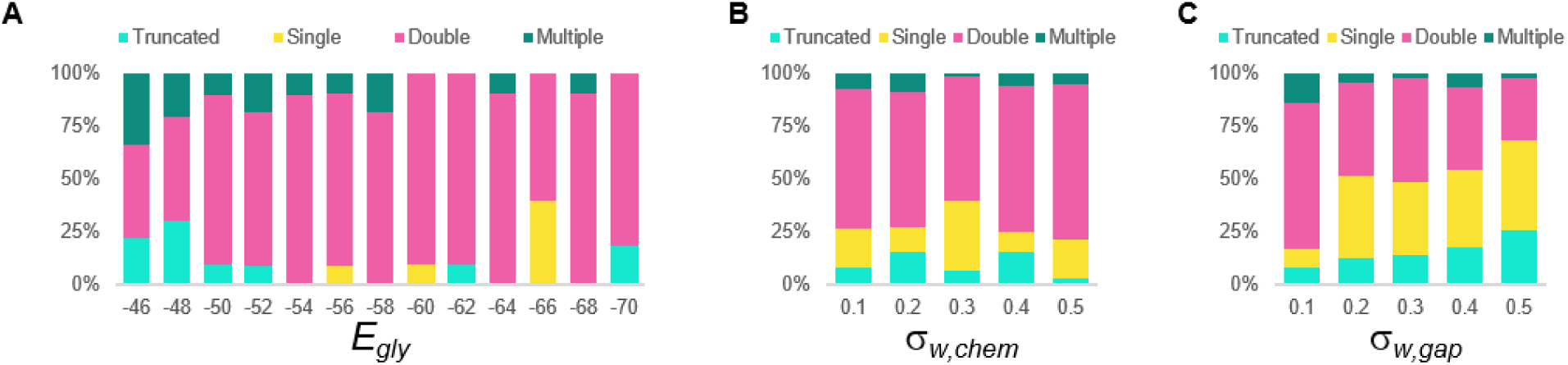
Sensitivity testing of the double coiling model for the glycinergic reversal potential (*E_gly_*), weights of chemical synapses (*σ_w, chem_*), and weights of gap junctions (*σ_w, gap_*). Sensitivity testing showing proportions of single, double, multiple, and truncated coiling events during ten 100,000 ms runs for each value tested.

**Figure 5 - figure supplement 1.**
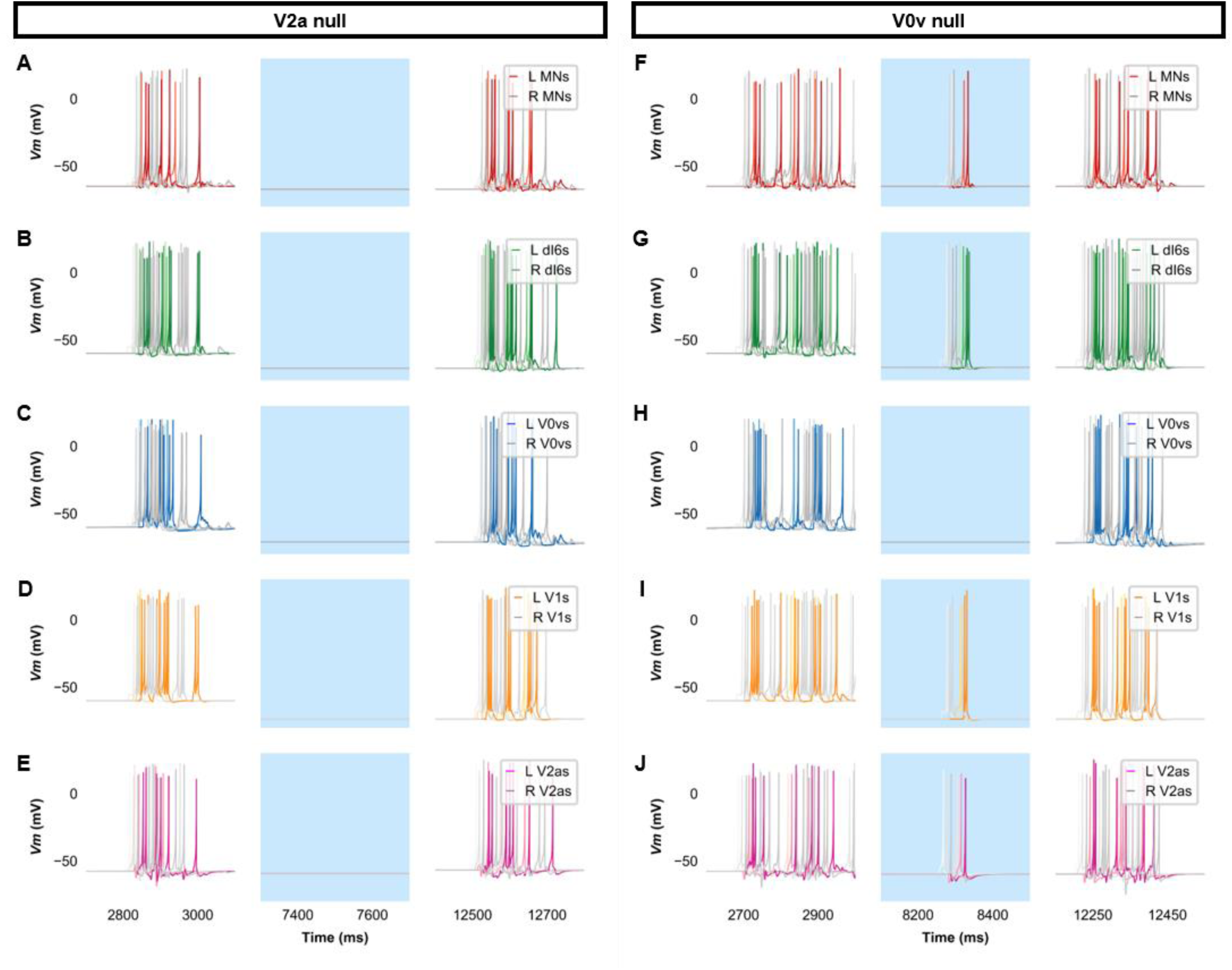
Membrane potential (*Vm*) of spinal neurons during simulations of beat-and-glide swimming where excitatory neurons were silenced. Simulations consisted of three 5,000 ms epochs. In the middle epoch, silencing of targeted spinal neurons was achieved by removing all synaptic and external currents from the targeted population. Synaptic and external currents were restored in the last epoch. (**A*-*E**) Simulations where V2as were silenced and (**F-J**), where V0vs were silenced in the middle epoch. The *Vm* of a rostral (lightest), middle, and caudal (darkest) neuron is shown. L: left, R: right.

**Figure 6 - figure supplement 1.**
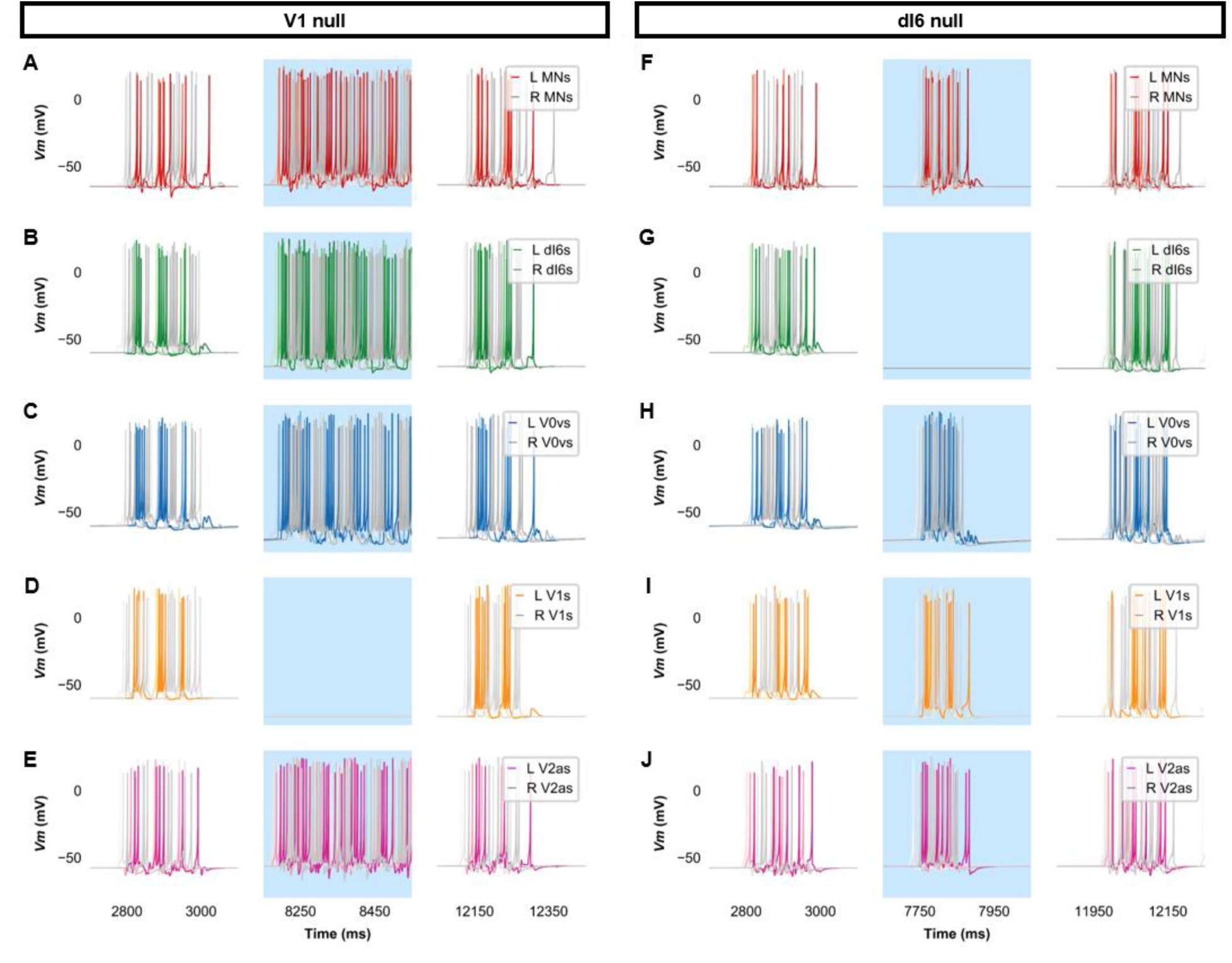
Membrane potential (*Vm*) of spinal neurons during simulations of beat-and-glide swimming where inhibitory neurons were silenced. Simulations consisted of three 5,000 ms epochs. In the middle epoch, silencing of targeted spinal neurons was achieved by removing all synaptic and external currents from the targeted population. Synaptic and external currents were restored in the last epoch. (**A*-*E**) Simulations where V1s were silenced and (**F-J**), where dI6s were silenced in the middle epoch. The *Vm* of a rostral (lightest), middle, and caudal (darkest) neuron is shown. L: left, R: right.

**Figure 6 - figure supplement 2.**
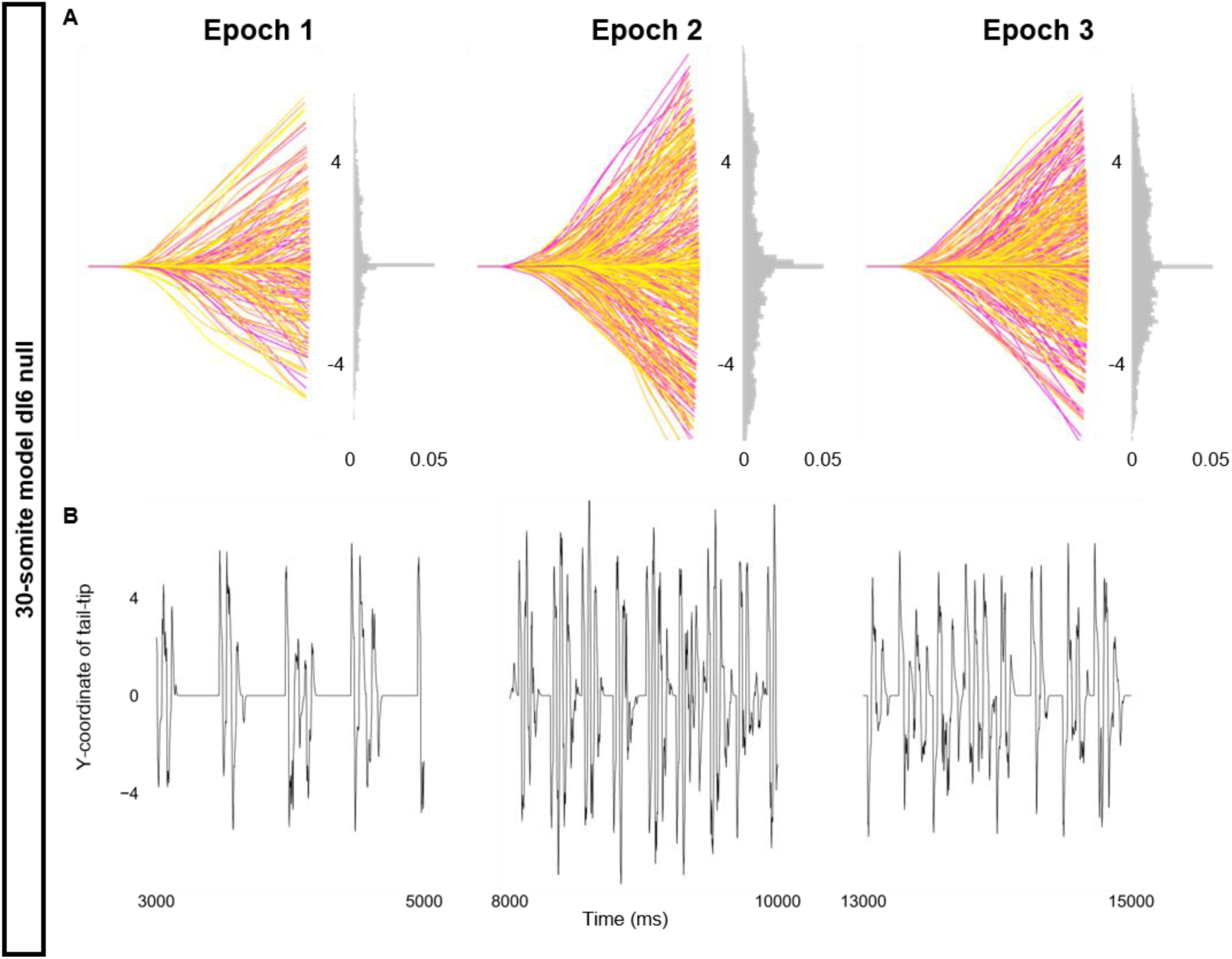
Altered kinematics during silencing of dI6 neurons. Simulation of a 30-somite beat-and-glide swimming model consisted of three 5,000 ms epochs. In the middle epoch, silencing of dI6s was achieved by removing all synaptic and external currents from the targeted population. Synaptic and external currents were restored in the last epoch. (**A**) Representative body midlines are shown for each epoch along with a probability density histogram of the y-coordinate of the terminal somite during each epoch. The histograms are truncated at 0.05 as there were many points at y = 0 during inter-episode intervals. The magenta to yellow color coding represents the progression through each epoch. (**B**) Y-coordinate of the tail tip during the last 2,000 ms of each epoch. Details of the 30-somite model are described in **Figure 8 - figure supplement 1.**

**Figure 7 - figure supplement 1.**
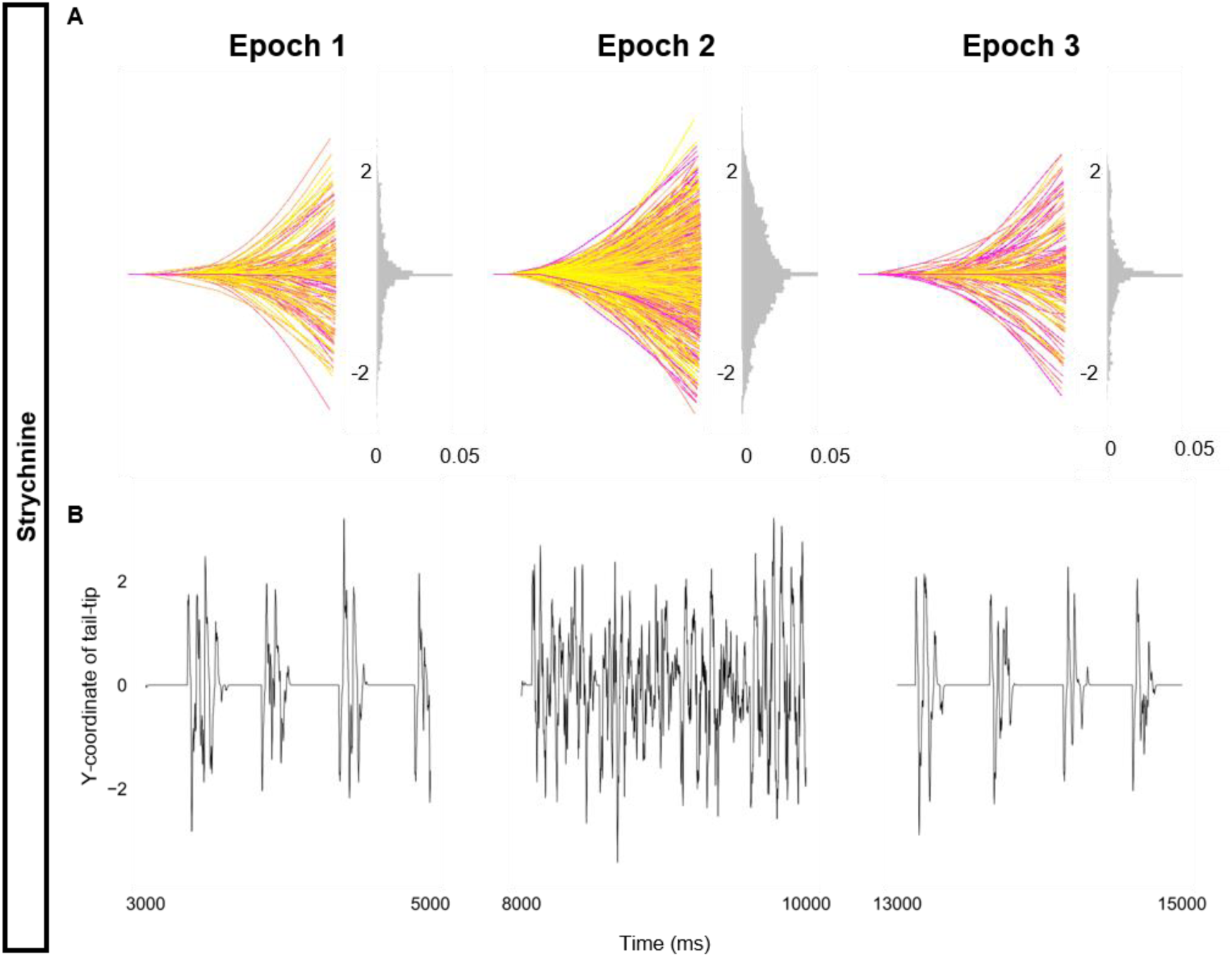
Altered kinematics during strychnine. Simulation of the base beat-and-glide swimming model consisted of three 5,000 ms epochs. In the middle epoch, all glycinergic currents were blocked. Glycinergic transmission was restored in the last epoch. (**A**) Representative body midlines are shown for each epoch along with a probability density histogram of the y-coordinate of the terminal somite during each epoch. The histograms are truncated at 0.05 as there were many points at y = 0 during inter-episode intervals. The magenta to yellow color coding represents the progression through each epoch. (**B**) Y-coordinate of the tail tip during the last 2,000 ms of each epoch.

**Figure 8 - figure supplement 1.**
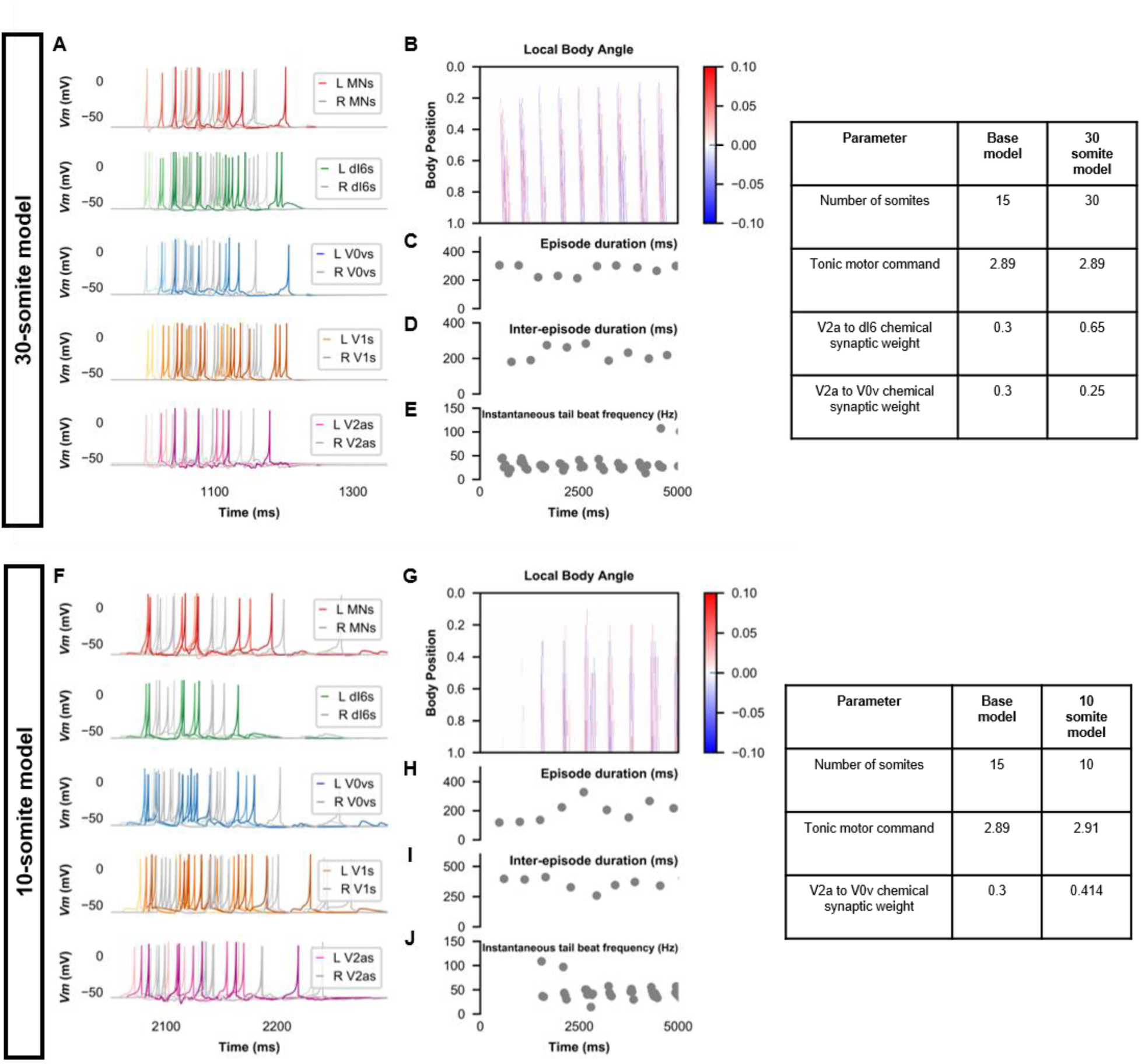
Beat-and-glide swimming model with different number of somites. (**A**, **F**) Membrane potential (*Vm*) of spinal neurons during a beat-and-glide swimming simulation. The *Vm* of a rostral (lightest), middle, and caudal (darkest) neuron is shown. L: left, R: right. (**B**, **G)** Heat-map of local body angle, (**C, H**) episode duration, (**D, I**) inter-episode interval, and (**E**, **J**) instantaneous tail beat frequency during the same simulations as **A** and **F**, respectively.

**Figure 8 - figure supplement 2.**
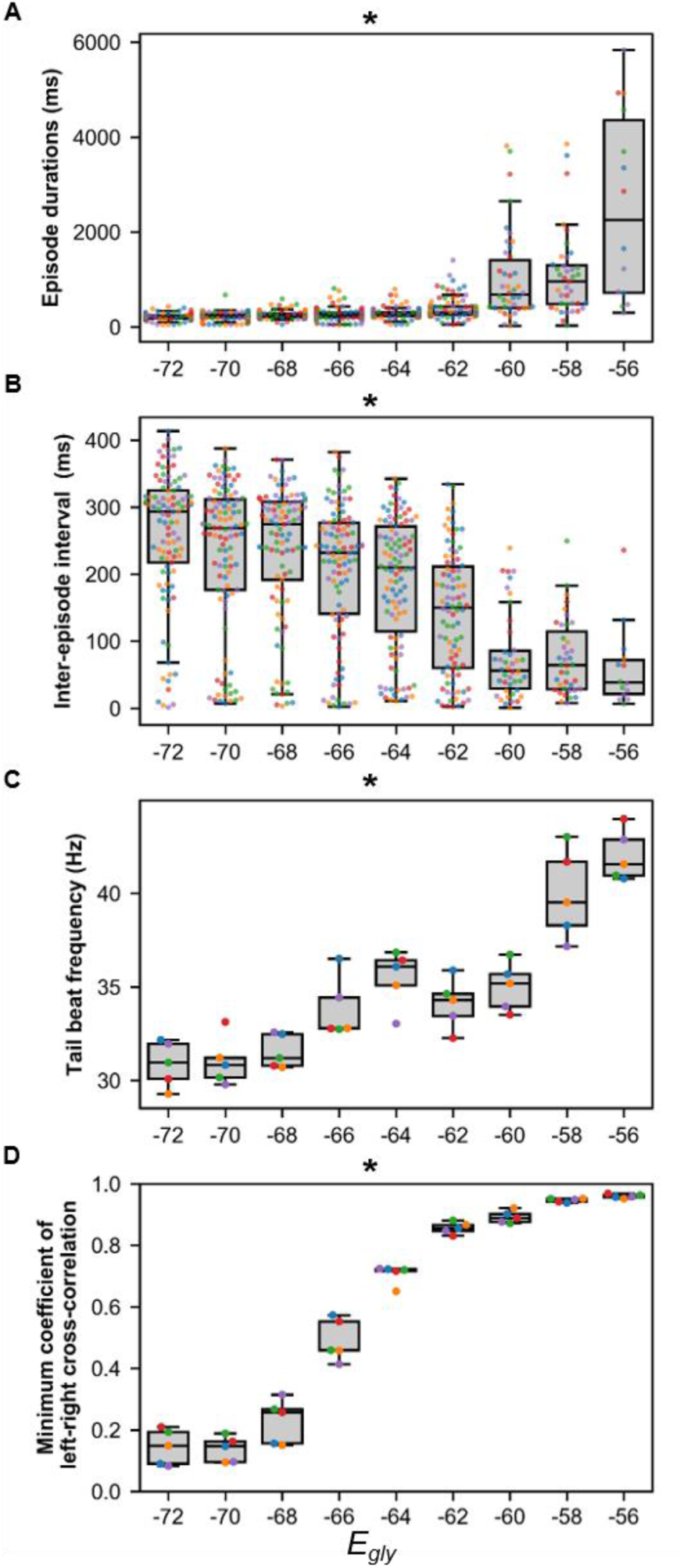
Sensitivity of beat-and-glide swimming to variability in glycinergic reversal potential (*E_gly_*). Five 10,000-ms long simulations were run for each value of *E_gly_*. (**A**) Episode duration, (**B**) inter-episode intervals, and (**C**) average tail beat frequency during each swimming episode. (**D**) The minimum coefficient of the cross-correlation of left and right muscle was calculated at each *E_gly_*. The minimum coefficient was taken between -10 and 10 ms time delays. Asterisks denote significant differences detected using a one-factor ANOVA test. Each run is color coded. ***Statistics:*** (**A**) *F*_8,681_ = 74.9, p = 2.7 x 10^-88^. (**B**) *F*_8,681_ = 32.6, p = 1.5 x 10^- 43^. (**C**) *F*_8,36_ = 22.9, p = 6.0 x 10^-12^. (**D**) *F*_8,36_ = 327.8, p = 3.0 x 10^-31^. P-values for t-tests are found in ***Figure 8* – *source data 1*.**

Figure 2 - *video 1* Single coiling model

Figure 2 - *video 2* Truncated coils

Figure 3 - *video 1* Double coiling model

Figure 3 - *video 2* Glutamate null double coiling model

Figure 3 - *video 3* Overexcited V0v double coiling model

Figure 3 - *video 4* Glycine null double coiling model

Figure 3 - *video 5* 30-somite double coiling model

Figure 4 - *video 1* Beat-and-glide model

Figure 5 - *video 1* V2a knockout beat-and-glide model

Figure 5 - *video 2* V0v knockout beat-and-glide model

Figure 6 - *video 1* V1 knockout beat-and-glide model

Figure 6 - *video 2 dI6* knockout beat-and-glide model

Figure 7 - *video 1* Glycine null beat-and-glide model

Figure 8 - *video 1* Beat-and-glide with bursting V2a model

Figure 8 - *video 2* Swimming model with only tonic neurons

Figure 8 - *video 3* 30-somite beat-and-glide model

Figure 2 - source data 1

Figure 5 - source data 1

Figure 6 - source data 1

Figure 7 - source data 1

Figure 8 - source data 1

Figure 9 - source data 1

Figure 10 - source data 1

